# Novel Fluorescent and Photoconvertible Fusions Reveal Dorsal Activator Dynamics

**DOI:** 10.1101/2025.05.12.653543

**Authors:** Meghan A. Turner, Nicholas M. Gravina, Bruno Moretti, Sadia Dima, Gabriella Martini, Greg Reeves, Hernan G. Garcia

**Author notes:** **For correspondence:** (HGG). These contributed equally to this work (Nicholas M. Gravina, Bruno Moretti, Sadia Dima).

## Abstract

Over the last two decades, new *in vivo* and *in cellulo* imaging technologies have uncovered the inherently dynamic nature of transcriptional regulation in embryonic development and, in particular, in the fruit fly *D. melanogaster*. These technologies have made it possible to characterize the subnuclear and single-molecule dynamics of transcription factors. However, a lack of appropriate fluorescent protein fusions has, until now, limited these studies to only a few of the dozens of important transcription factors in the fruit fly gene regulatory network dictating early development. Here, we report the creation of four new fluorescent protein fusions to Dorsal, a member of the NF-*κ*B/Rel family that initiates dorsal-ventral patterning. We generated and characterized two bright fluorescent protein fusions for Dorsal, meGFP and mNeonGreen, and two photoconvertible fluorescent protein fusions, mEos4a and Dendra2. We show that removal of the DsRed2 cassette commonly used to mark the CRISPR integration restores endogenous Dorsal mRNA and protein levels and enables the fusion allele to rescue a *dorsal* null allele, meeting the gold standard for endogenous function of a tagged protein in a fruit fly. We then demonstrate that our bright fluorescent protein fusions can be used to dissect the spatiotemporal dynamics of stable Dorsal clusters that traverse the nucleoplasm and uncovered that these clusters preferentially interact with active sites of Dorsal-modulated transcription. We further demonstrate that our photoconvertible fluorescent protein fusions make it possible to detect individual molecules of Dorsal in the nuclei of developing embryos. These new fluorescent protein fusions constitute a valuable resource for the community to elucidate the role of Dorsal activator dynamics in dictating fruit fly early embryonic development.

## 1 Introduction

Over the last two decades, new *in vivo* and *in cellulo* microscopy technologies have made it possible to uncover the dynamics of regulatory transcription factors as they interact with transcription sites to activate or repress gene expression in the context of embryonic development (***Wagh et al., 2023***; ***Boka et al., 2021***; ***Lu and Lionnet, 2021***). The emerging picture is one where activators or repressors only transiently occupy their target binding sites at enhancers (***Mir et al., 2017, 2018a***; ***Lu and Lionnet, 2021***; ***Donovan et al., 2019***)—and sometimes act in the context of spatially localized hubs or clusters (***Mir et al., 2017, 2018a***; ***Sabari et al., 2018***; ***Wei et al., 2020***; ***Kawasaki and Fukaya, 2023***)—to regulate the stochastic transcription process underpinned by transcriptional bursting (***Rodriguez and Larson, 2020***; ***Lammers et al., 2020b***; ***Leyes Porello et al., 2023***; ***Meeussen and Lenstra, 2024***).

These discoveries have been partially fueled by an ever-increasing palette of fluorescent proteins, which are fused to transcription factors to enable direct measurements of their real-time dynamics. This palette now includes fluorescent proteins that are suitable for a wide range of live imaging experiments: brighter and more photostable fluorescent proteins (meGFP, mClover3, mNeonGreen, mStayGold) enable longer-term imaging, and photoactivatable (PA-GFP) and photo-convertible (mEos3.2, Dendra2) fluorescent proteins enable superresolution and single-molecule imaging (all reviewed in ***Rodriguez et al. (2017***)).

Yet, despite this ever-growing toolbox of fluorescent proteins, it is time-consuming and challenging to fuse newly engineered fluorescent proteins to a protein of interest in a manner that preserves that protein’s endogenous functionality (***Chen et al., 2011***; ***Cranfill et al., 2016***). For example, it has proven particularly difficult to generate fluorescent protein fusions for early transcription factors (TFs) in the developing fruit fly (*Drosophila melanogaster*) embryo, such as Bicoid and Dorsal (***Reeves et al., 2012***; ***Singh et al., 2022***).

Dorsal, a transcriptional activator belonging to the NF-*κ*B/Rel family (***Hong et al., 2008***; ***Gilmore, 2006***), initiates fruit fly embryonic dorsal-ventral patterning via a maternally deposited concentration gradient (***Hong et al., 2008***; ***Gilmore, 2006***). Despite its crucial role in the developmental cascade of the early fruit fly embryo, studies of Dorsal dynamics have been limited by the availability of functional fluorescent protein fusions. Only a single fluorescent protein fusion, Dorsal-mVenus, meets the gold standard for maintaining endogenous Dorsal activator activity: a single copy of a Dorsal-mVenus transgene allele complements (or, “rescues” the development of) a Dorsal null allele (e.g. *<u>dl[1]</u>*). Subsequently, ***Alamos et al. (2023***) successfully generated a Dorsal-mVenus CRISPR knock-in allele—using the same combination of linker and fluorescent protein as in the transgene by ***Reeves et al. (2012***)—that also rescues the *dl[1]* null allele.

While Dorsal-mVenus has proven exceedingly useful in measuring the nuclear levels and dorso-ventral gradient of Dorsal in the embryo (***Reeves et al., 2012***; ***Alamos et al., 2023***; ***Pimmett et al., 2024***), we sought to answer outstanding questions about the sub-nuclear dynamics of Dorsal, including measuring the activity of subnuclear clusters and the binding dynamics of single Dorsal molecules. These specific experimental goals required a fluorescent protein more photostable than mVenus, as well as a photoconvertible fluorescent protein.

Here, we describe an expansion of the fluorescent protein palette for endogenous Dorsal, featuring four new fluorescent protein CRISPR knock-in fusions: the brighter and more photostable meGFP (***Cormack et al., 1996***; ***Zacharias et al., 2002***) and mNeonGreen (***McKinney et al., 2009***), and the photoconvertible mEos4a (***Kopek et al., 2017***; ***Paez-Segala et al., 2015***) and Dendra2 (***Gurskaya et al., 2006***). All four fusions produce viable progeny from females homozygous for the Dorsal-FP allele, and we show that removing the DsRed marker cassette enables our Dorsal-mNeonGreen allele to rescue a Dorsal null allele. Thus, these new fusions are suitable for studying the endogenous dynamics of Dorsal.

We demonstrate the potential of these brighter and more photostable meGFP and mNeon-Green fusions to study recently discovered clusters of Dorsal concentration that have been suggested to play a role in the regulation of Dorsal target genes (***Yamada et al., 2019***). Our fusions make it possible to track the dynamics of these clusters, revealing that Dorsal clusters tend to be in closer proximity to target genes than non-target genes—a phenomenon that we explore more deeply in a pair of accompanying papers (***Fallacaro et al., 2025***; ***Dima et al., 2025***). Additionally, we demonstrate how the photoconvertible mEos4a and Dendra2 fusions enabled us to track single molecules of Dorsal binding to DNA for the first time, finding that Dorsal spends only a few seconds bound to the DNA, an observation consistent with the binding times of several other transcription factor in the fruit fly and beyond (***Lammers et al., 2020b***; ***Lu and Lionnet, 2021***). These two experiments, only made possible by the new Dorsal fusion alleles, help increase our understanding of the dynamic process of transcriptional activation and demonstrate that the four Dorsal fusions presented in this paper will constitute a valuable resource for the community.

## 2 Results

### 2.1 Generation of novel Dorsal fusions

We fused two fluorescent proteins, meGFP and mNeonGreen (***McKinney et al., 2009***), and four photoconvertible proteins, Dendra2 (***Gurskaya et al., 2006***), mEos3.2 (***Zhang et al., 2012***), mEos4a, and mEos4b (***Paez-Segala et al., 2015***; ***Kopek et al., 2017***), in-frame to the C-terminus of the Dorsal protein via a 6xGlycine (6G) linker using an existing CRISPR/Cas9 protocol (***Gratz et al. (2015***); ***Alamos et al. (2023***); *Methods* ***Section 4.1*** and ***Table S1***). The knock-in was marked by a 3xP3-DsRed2-SV40polA cassette, which drives the expression of the fluorescent protein DsRed in the adult eyes and ocelli, allowing for rapid screening of successfully transformed adults. This DsRed cassette, as we will refer to it from now on, was flanked by a pair of 3’ and a 5’ PiggyBac transposon sites to allow for scarless removal of DsRed. The DsRed cassette was placed downstream of the fluorescent protein’s stop codon (***Gratz et al., 2015***) and upstream of the endogenous 3’ untranslated region (3’UTR). We characterized several successful (as indicated by DsRed+ adults), independent integrations of each CRISPR knock-in *dorsal* (*dl*) allele.

The introduction of the fluorophore sequence can interfere with the regulation, production, or function of the protein to which it is fused, resulting in impaired downstream functions. As an initial assessment of the function of these fusions, we tested whether females homozygous for the Dorsal fusion alleles were fertile and able to generate viable progeny. In the case of a maternally-deposited transcription factor like Dorsal, the ability for females homozygous for the fusion alleles to produce viable progeny indicates that the developmental functions of Dorsal are intact. Females homozygous for the *dl-6G-mEos4b-DsRed* and *dl-6G-mEos3.2-DsRed* alleles were not fertile, generating no viable progeny at room temperature (approximately 22°C). This outcome indicates that the maternally deposited copies of these Dorsal fusion proteins do not function well enough to drive development (***Table 1***). As a result, we did not proceed with any further characterization of the mEos4b nor mEos3.2 lines.

**Table 1.** Summary of the *dl* alleles characterized in this study. For alleles with more than one entry, two unique CRISPR/Cas9 integrations were made into stable lines (labelled as e.g. “line 1” and “line 8”) and characterized. Embryo hatch rate was quantified separately for each integration; all other metrics were combined between the two integrations of the same allele. Linker protein sequences: 6G (GGGGGG); LL (LongLinker; SGDSGVYKTRAQASNSAVDGTAGPGSTGSS); 6G-10GS (GGGGGGGGGGSGSGGS); 6G-helix (GGGGGGMSKGEEL; MSKGEEL is the N-terminal helix from mVenus). ^*a*^The number of DsRed+ adults divided by the total number of injected embryos that survived to adulthood. “Unsuccessful” indicates no injected embryos survived to adulthood. ^*b*^females homozygous for the CRISPR allele (i.e. females deposit no wild-type Dorsal into the embryo) produce viable pupae. ^*c*^Fraction of embryos laid by homozygous females that hatch after 36 hours. ^*d*^One copy of the CRISPR allele complements (rescues function) a *dl* null allele. Either *dl[1]* or *dl[4]* were used. ^*^Not tested

| Allele | CRISPR efficiency<sup>a</sup> | Homozyg. females produce pupae at 22°C<sup>b</sup> | Homozyg. females produce pupae at 25°C<sup>b</sup> | Embryo hatch rate<sup>c</sup> | Rescues *dl* null allele<sup>d</sup> |
| --- | --- | --- | --- | --- | --- |
| *dl* (wild-type) | N/A | yes | yes | 0.85 | yes |
| *dl-6G-mVenus-DsRed* | unknown | yes | yes | 0.75 | yes |
| *dl-6G-Dendra2-DsRed* (line 1) | 0.05 | yes | no | 0.55 | —* |
| *dl-6G-Dendra2-DsRed* (line 2) |  |  |  | 0.60 | — |
| *dl-6G-meGFP-DsRed* (line 1) | 0.10 | yes | no | 0.05 | — |
| *dl-6G-meGFP-DsRed* (line 2) |  |  |  | 0.20 | — |
| *dl-6G-mEos4a-DsRed* (line 1) | 0.05 | yes | no | 0.15 | — |
| *dl-6G-mEos4a-DsRed* (line 8) |  |  |  | 0.20 | — |
| *dl-6G-mNeonGreen-DsRed* (line 2) | 0.06 | yes | no | 0.02 | no |
| *dl-6G-mNeonGreen-DsRed* (line 6) |  |  |  | 0.05 | no |
| *dl-LL-mNeonGreen-DsRed* | 0.60 | yes | no | 0.01 | — |
| *dl-6G-mNeonGreen-ΔDsRed* | N/A | yes | — | — | yes |
| *dl-6G-mEos4b-DsRed* | 0.14 | no | N/A | 0.00 | — |
| *dl-6G-mEos3.2-DsRed* | 0.10 | no | N/A | 0.00 | — |
| *dl-6G-10GS-mEos3.2-DsRed* | unknown | no | N/A | N/A | N/A |
| *dl-6G-helix-mEos3.2-DsRed* | unknown | no | N/A | N/A | N/A |
| *dl-LL-mEos3.2-DsRed* | unknown | no | N/A | N/A | N/A |

In contrast, females homozygous for the *dl-6G-meGFP-DsRed, dl-6G-mNeonGreen-DsRed, dl-6G-mEos4a-DsRed*, and *dl-6G-Dendra2-DsRed* alleles yielded viable pupae at room temperature. These results indicate that these Dorsal fusion proteins maintain some level of normal function during development. However, very few pupae were produced by females homozygous for these four alleles. Only females homozygous for the *dl-6G-Dendra2-DsRed* allele produced sufficient progeny to be maintained as a stable, homozygotic line; females homozygous for the other three alleles—*dl-6G-meGFP-DsRed, dl-6G-mNeonGreen-DsRed*, and *dl-6G-mEos4a-DsRed*—produced too few progeny to be maintained as stable, homozygotic line. Only the *dl-6G-Dendra2-DsRed* allele produced sufficient progeny to be maintained as a stable, homozygotic line. Additionally, when females homozygous for these four alleles were kept at elevated temperatures (25°C), they no longer yielded pupae, suggesting that these fusion alleles compromise the robustness of the function of Dorsal during development in the face of temperature changes.

As a more quantitative measure of fly line viability, we measured and compared the embryo hatch rate across these four homozygous alleles (***Figure 1A***). We counted the percentage of embryos, laid by females homozygous for each allele, that successfully hatched into larvae after 36 hours (*Methods* ***Section 4.4***). We compared these hatch rates to the wild-type (untagged) *dl* allele (*yw* ; *+ ;+*, hereafter *yw*) and the *dl-6G-mVenus-DsRed* allele (***Alamos et al., 2023***). Embryos from wild-type *dl* females had the highest hatch rate, 85%, followed closely by embryos from *dl-6G-mVenus-DsRed* females, which hatched at a 75% rate. None of the embryos from females with the new Dorsal fusion alleles exceeded the hatch rate of the *dl-6G-mVenus-DsRed* allele. *dl-6G-Dendra2-DsRed* had the highest hatch rate of the new alleles (55% and 60%, for two independent integrations), followed by *dl-6G-mEos4a-DsRed* (15% and 20%) and *dl-6G-meGFP-DsRed* (5% and 20%). *dl-6G-mNeonGreen-DsRed* exhibited the lowest embryo hatch rates (2% and 5%).

**Figure 1.**
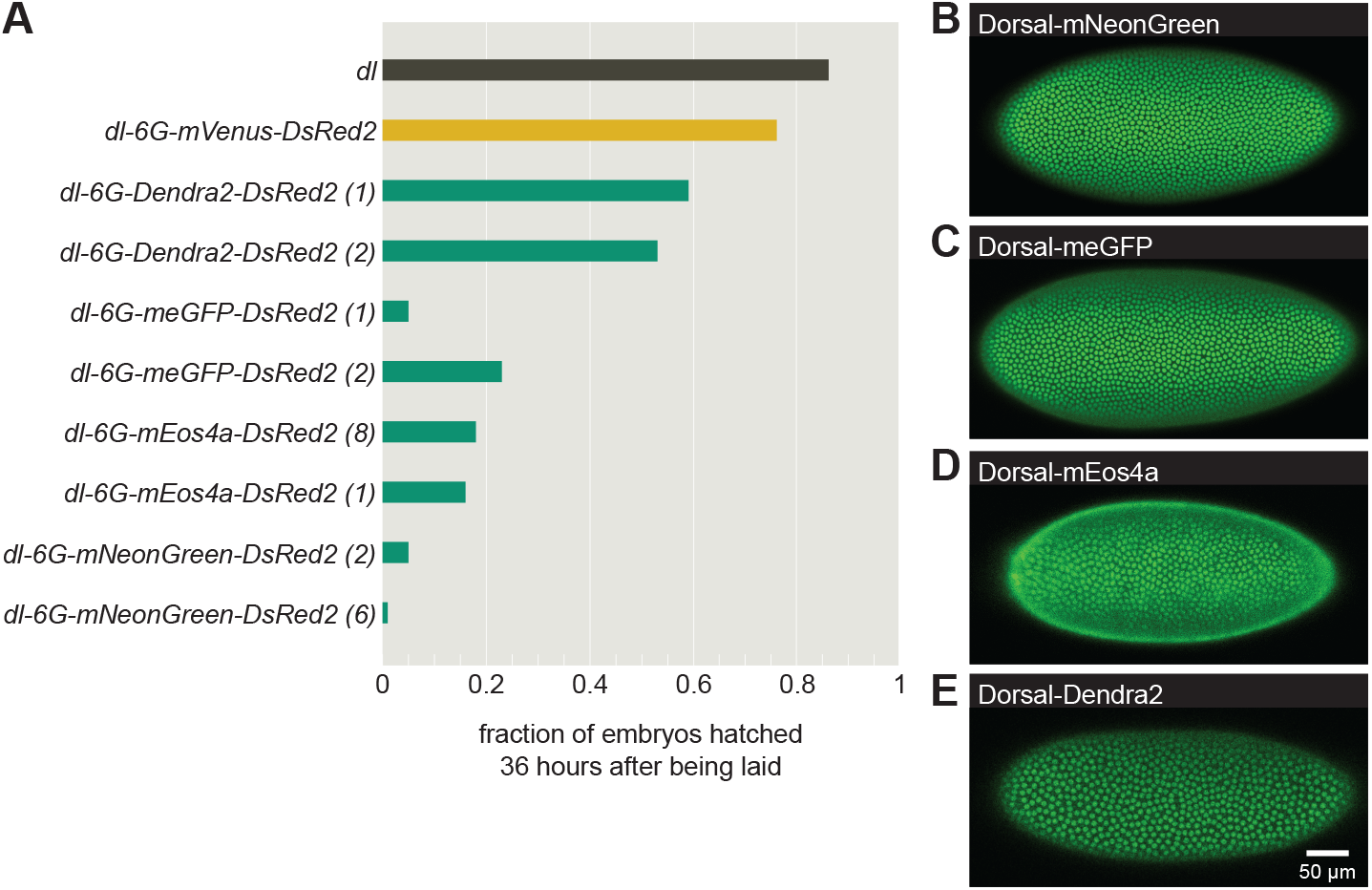
Generation and characterization of novel Dorsal fusion proteins. **(A)** Embryo hatch rates for embryos laid by females homozygous for various Dorsal fusion alleles, quantified as fraction of embryos that had hatched 36 hours after being laid. Embryo hatch rates for an untagged *dl* allele (from *yw* flies) served as a control. Hatch rates for each allele can also be found in ***Table 1*. (B-E)** Images of our homozygous-viable Dorsal fusion lines taken on laser scanning confocal microscope during later nuclear cycle 14 prior to gastrulation: **(B)** Dorsal-mNeonGreen, **(C)** Dorsal-meGFP, **(D)** Dorsal-mEos4a, and **(E)** Dorsal-Dendra2.

To further characterize these fusions, we performed live imaging to assess the resulting Dorsal protein expression pattern. All four alleles drive a Dorsal gradient that qualitatively matches the expected endogenous expression pattern of Dorsal protein: high nuclear Dorsal levels along the ventral midline decreasing to nuclear exclusion of Dorsal towards the dorsal side of the embryo ***Figure 1B-E***. Thus, our results demonstrated, at least qualitatively, that the maternal Dorsal protein was being translated and imported to the nuclei as expected.

Despite the qualitative agreement between the endogenous Dorsal gradient and the gradient resulting from our Dorsal fusions, the poor viability of our new fusion fly lines (***Table 1***) led us to question whether these lines were faithful reporters of endogenous Dorsal function. Specifically, we hypothesized that the poor viability of our CRISPR *dl* alleles could have three causes: off-target CRISPR/Cas9-induced mutations, interference from the protein sequence linker, and/or interference from the presence of the DsRed cassette in the 3’UTR.

### 2.2 Removal of DsRed cassette restores embryo viability of Dorsal fusion fly lines

We investigated and corrected for the three potential causes of the poor embryo viability in our homozygous fusion allele fly lines, as hypothesized in the previous section. First, we removed off-target CRISPR/Cas9-induced mutations in other essential genes via out-crossing. Such mutations could lead to non-viability, without implicating maternal Dorsal function. Out-crossing for six to eight generations did not improve embryo viability (Supplemental Information ***Section S1*** ; ***Figure S1***).

Second, we altered the linker sequence between Dorsal and the fluorescent protein fusion. The specific sequence of certain linkers may interfere with the regulation, folding, or function of maternal Dorsal more than the sequence of others. We were unable to identify an alternative linker sequence that improved the embryo viability of these fly lines (Supplemental Information ***Section S2*** ; ***Table 1***).

Third, we assessed the effect of the DsRed marker cassette in the 3’UTR. The DsRed cassette is located between the stop codon of the fluorescent protein and the start of the endogenous 3’ untranslated region (3’UTR) sequence. While this position does not alter the Dorsal protein coding sequence, it does modify the Dorsal mRNA sequence, particularly the position and sequence of its 3’UTR. 3’UTRs play a significant role in mRNA stability, mRNA localization, regulation of translation, and protein-protein interactions (***Szostak and Gebauer, 2013***; ***Buxbaum et al., 2015***; ***Andreassi and Riccio, 2009***; ***Kuersten and Goodwin, 2003***; ***Mayr, 2019***). The presence of the DsRed cassette in the 3’UTR could interfere with any of these important molecular functions, leading to altered Dorsal protein expression levels and impacting downstream target genes.

To assess the effect of the DsRed cassette on viability, we removed this sequence from the line carrying the *dl-6G-mNeonGreen-DsRed* allele using scarless removal via the PiggyBac transposase (***Nyberg and Carthew, 2022***). We then quantified and compared the viability of the allele with and without the DsRed marker cassette. In contrast to embryos from females with two copies the *dl-6G-mNeonGreen-DsRed* allele, embryos from females with two copies of the *dl-6G-mNeonGreen-* Δ*DsRed* allele were viable and could produce a stable stock population. Given this improvement in embryo viability, we then tested our *dl-6G-mNeonGreen-*Δ*DsRed* allele for its ability to maintain embryo viability as a single copy. The ability of a transgenic *dl* allele to complement a mutant null *dl* allele–here, *dl[4]*—is considered the gold standard for demonstrating normal protein function. We found that embryos from *dl-6G-mNeonGreen-*Δ*DsRed* / *dl[4]* females were viable, but embyros from *dl-6G-mNeonGreen-DsRed* / *dl[4]* females were not. Thus, our results indicate that the interruption in the 3’UTR reduces embryo viability.

### 2.3 Changes in the *dorsal* 3’UTR modulate mRNA levels and nuclear protein concentration

To identify the molecular cause of this reduced embryo viability, we measured and compared the *dl* mRNA levels, Dorsal protein pattern, nuclear Dorsal protein concentrations, and Dorsal-target gene expression produced by the *dl-6G-mNeonGreen* allele with and without the DsRed cassette present, hereafter referred to as *DsRed* and Δ*DsRed*, respectively.

First, we quantified the effect of the DsRed cassette on *dl* mRNA production by measuring mRNA levels using quantitative real-time PCR (qPCR) on embryos collected from homozygous females of the *DsRed* and the Δ*DsRed* lines. We determined the relative mRNA levels with respect to *yw* embryos carrying a wild-type *dl* allele (***Figure 2A***). The ratio of *dl-6G-mNeonGreen-DsRed* to wild-type *dl* mRNA levels was 0.5±0.1 (mean ± standard error of the mean), indicating that the presence of the DsRed cassette reduced mRNA levels by half. In contrast, the ratio between *dl-6G-mNeonGreen-* Δ*DsRed* and wild-type *dl* was found to be 1.0±0.2, indicating that the scarless removal of the DsRed cassette restored *dl* mRNA levels to wild-type. As a result, we concluded that the significant reduction in mRNA levels due to the presence of the DsRed cassette in the 3’UTR of the *dl* allele was a likely cause of the reduced viability of the fly lines carrying the *dl-6G-mNeonGreen-DsRed* allele.

**Figure 2.**
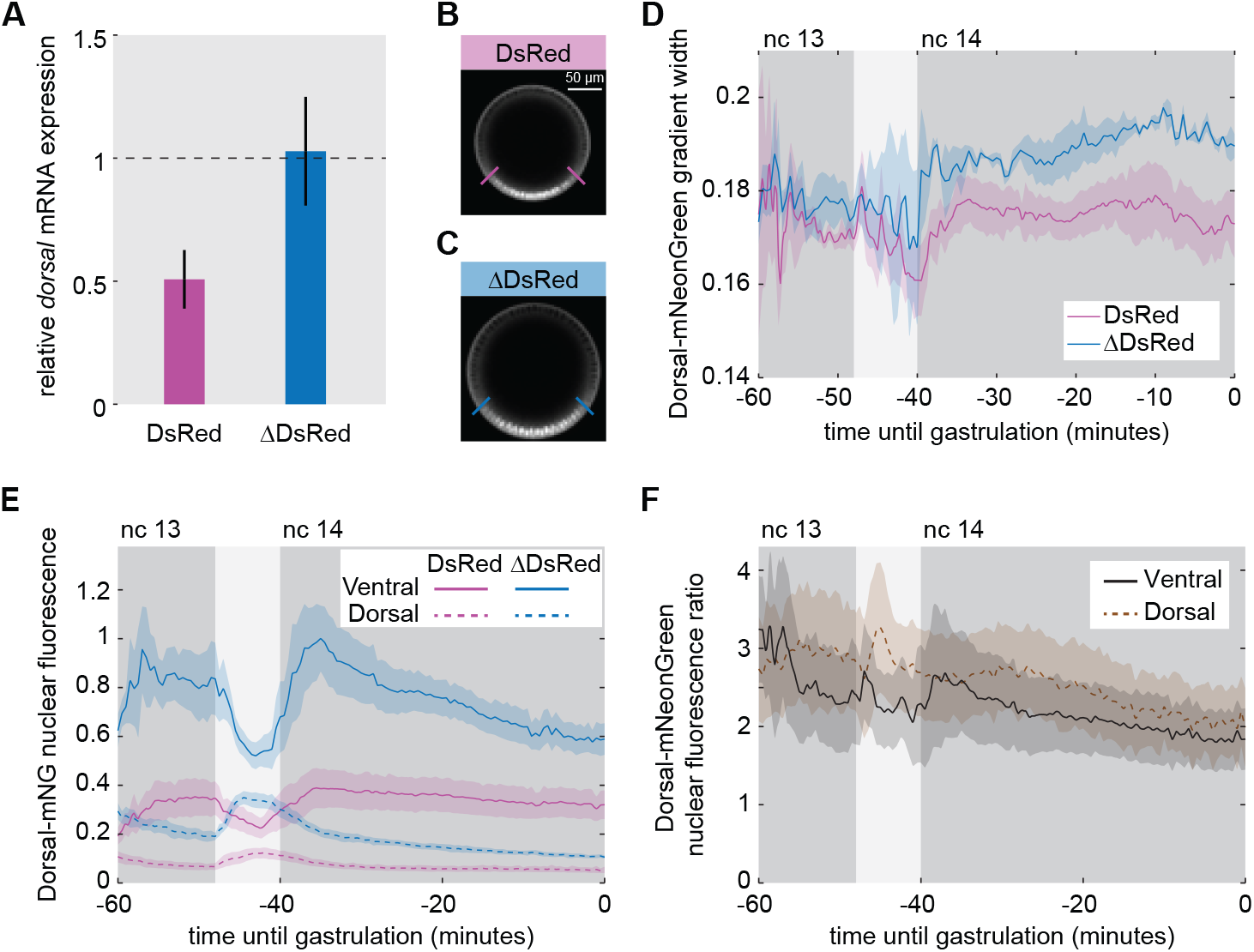
Effect of the presence of DsRed in the *dl* 3’-UTR. **(A)** *dl* mRNA expression ratio, as measured by qPCR, for embryos from females carrying two copies of the *dl-6G-mNeonGreen-DsRed* (DsRed; pink) and the *dl-6G-mNeonGreen-*Δ*DsRed* (ΔDsRed; blue) alleles, relative to a wild-type *dl* allele. *dl-6G-mNeonGreen-DsRed*: 0.5±0.1; *dl-6G-mNeonGreen-*Δ*DsRed*: 1.0±0.2 (mean ± standard error of the mean). **(B-C)** Representative cross-section images of the Dorsal-mNeonGreen fluorescent protein gradient and measurement of gradient width, as indicated by the perpendicular colored lines, in **(B)** *dl-6G-mNeonGreen-DsRed* / His2Av-RFP embryos and **(C)** *dl-6G-mNeonGreen-*Δ*DsRed* / His2Av-RFP embryos. Scale bar is 50 *μ*m. **(D)** Width of the Dorsal-mNeonGreen gradient, as extracted from a Gaussian fit, in DsRed and ΔDsRed embryos during nuclear cycles 13 and 14 (dark regions) and the intervening mitosis (light region). **(E)** Quantification of the normalized Dorsal-mNeonGreen nuclear fluorescence in ventral (solid lines) and dorsal (dashed lines) nuclei for DsRed (pink) and ΔDsRed (blue) embryos during nuclear cycles 13 and 14. Line values are the mean of five DsRed embryos and three ΔDsRed embryos; shaded regions are SEM. **(F)** Ratio of Dorsal-mNeonGreen nuclear fluorescence in ΔDsRed to DsRed embryos in the ventral (solid black line) and dorsal (dashed, brown line) nuclei.

Second, to determine the downstream impact of these reduced mRNA levels on Dorsal protein levels and localization, we measured the Dorsal-mNeonGreen protein gradient along the dorsalventral axis during development. We imaged the cross-section of embryos from both heterozygous *dl-6G-mNeonGreen-DsRed* / *His2Av-RFP* (***Figure 2B***) and heterozygous *dl-6G-mNeonGreen-*Δ*DsRed* / *His2Av-RFP* females (***Figure 2C***), where the *His2Av-RFP* fluorescence signal was used for nuclear segmentation. The Dorsal-mNeonGreen protein gradient is qualitatively similar in the *DsRed* and Δ*DsRed* embryos, with high nuclear Dorsal-mNeonGreen levels in the ventral nuclei, lower nuclear Dorsal-mNeonGreen levels in dorsal nuclei, and negligible cytoplasmic Dorsal-mNeonGreen levels (***Figure 2B-C***). To quantify the Dorsal nuclear concentration gradient along the embryo, we fit a Gaussian function to the nuclear Dorsal-mNeonGreen signal across the full dorsal-ventral axis and determined its width and amplitude (*Methods* **??**; ***Liberman et al. (2009***); ***Reeves et al. (2012***)). Although the width of the Dorsal gradient was similar for *DsRed* and Δ*DsRed* embryos in nuclear cycle 13, we observed a slight difference in nuclear cycle 14 (***Figure 2D***). When we measured the amplitude of the Dorsal gradient, we found that the *DsRed* embryos had approximately half the nuclear Dorsal-mNeonGreen fluorescence in their ventral-most nuclei as compared to the Δ*DsRed* embryos (***Figure 2E-F***). Similarly, we found that the ventral nuclei in *DsRed* embryos contained a little more than half (approximately 53% in nuclear cycle 14) the absolute Dorsal-mNeonGreen protein concentration than the Δ*DsRed* embryos, as measured by Raster Image Correlation Spectroscopy (RICS) (Supplementary Information ***Section S3*** ; *Methods* **??**).

Finally, as expected, the reduced levels of nuclear Dorsal protein in the DsRed embryos led to altered mRNA expression patterns in downstream, Dorsal-regulated genes (***Figure S3***). We found that only 11% of the embryos had a wild-type expression pattern of the Dorsal-activated gene, *snail* in the ventral nuclei (***Figure S3B(i)***), with the remaining embryos exhibiting either a significantly reduced or entirely absent *sna* pattern (***Figure S3B(ii-iii)***). We posit that the aberrant expression of *snail* and other gene expression patterns led to the low percentage of hatched embryos of this and other transgenic lines (***Table 1***).

We conclude that the presence of the DsRed marker cassette in the Dorsal 3’UTR reduces the viability of the fly line by halving *dl* mRNA levels, which in turn leads to downstream reduction of Dorsal protein levels and severely altered gene expression of Dorsal-activated genes. We can restore the viability of these new fusion fly lines by scarless removal of the DsRed cassette from the 3’UTR.

### 2.4 Photostable fluorescent proteins uncover subnuclear Dorsal cluster dynamics and their proximity to sites of transcription

Recently, regions of high Dorsal concentration—clusters—were found in the vicinity of target genes (***Yamada et al., 2019***). These clusters were found to be correlated with an increase in the mean rate of transcription of Dorsal target genes. However, because these measurements were performed using fixed tissue techniques, how Dorsal cluster dynamics dictate transcriptional control remains unknown.

Using the existing Dorsal-mVenus CRISPR fusion (***Alamos et al., 2023***) line and live imaging on a confocal microscope, we uncovered relatively stable, submicron-sized clusters of high Dorsal concentration that move about the nucleoplasm (***Figure 3A***). These initial results confirmed that these clusters were not just an artifact of the original fixed embryo staining with anti-Dorsal antibody labeling that was used to discover them (***Yamada et al., 2019***).

**Figure 3.**
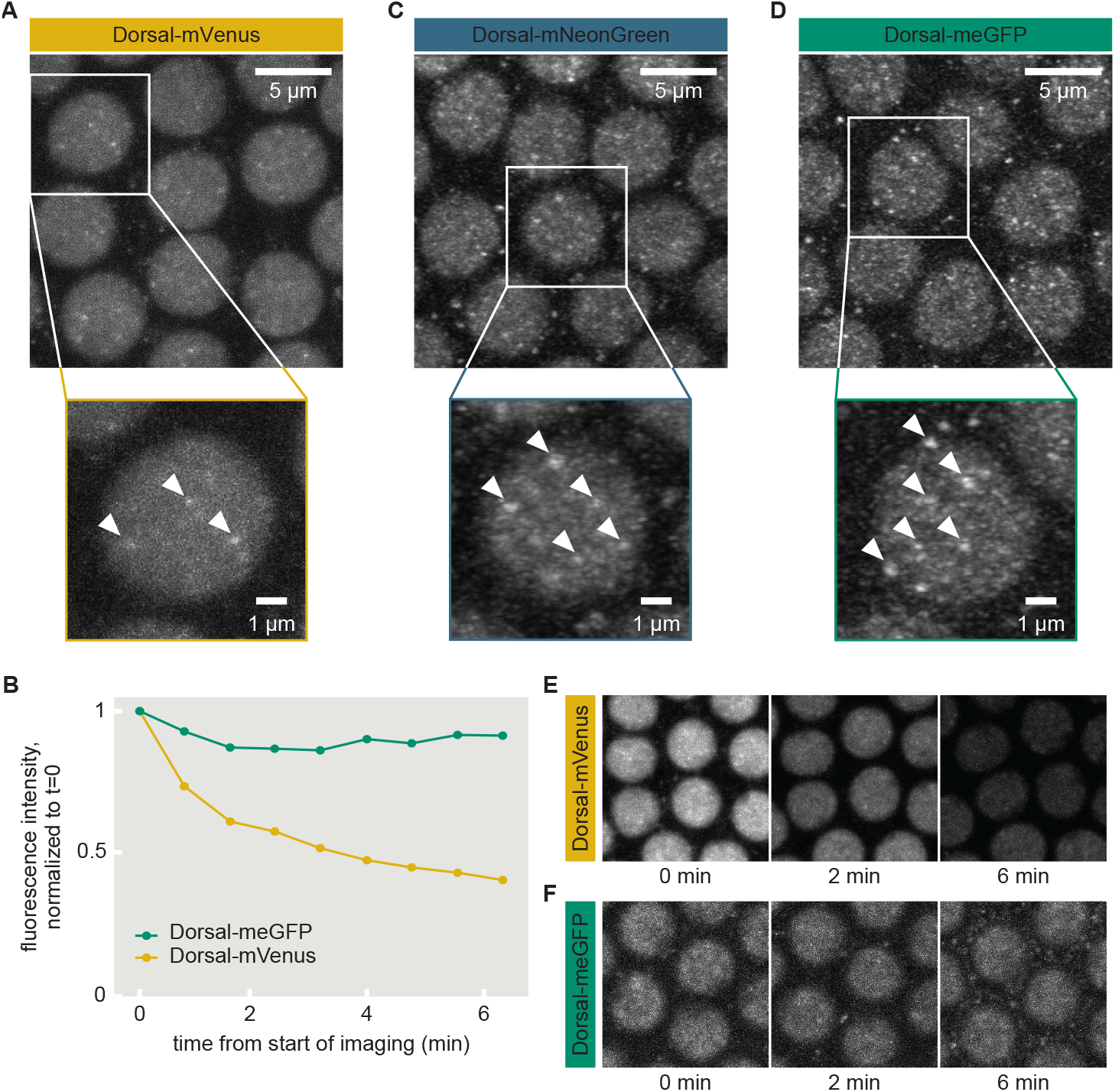
Dorsal-meGFP and Dorsal-mNeonGreen fusions enable live imaging of Dorsal clusters. **(A)** Snapshots from movies of Dorsal-mVenus revealing clusters of high concentration (white arrows). **(B)** Dorsal-mVenus clusters showed rapid bleaching as compared to the minimal bleaching exhibited by Dorsal-meGFP when imaged on the same microscope, under the same imaging conditions. **(C, D)** Dorsal clusters are clearly visible (white arrows) in the context of fusions to (C) mNeonGreen and (D) meGFP. **(E, F)** Representative snapshots of the field of view used to quantify the total fluorescence intensity for each time point quantified in (B) for (E) Dorsal-mVenus and (F) Dorsal-meGFP.

Yet, despite our ability to capture the clusters using the Dorsal-mVenus fusion, these clusters were only observable under a high spatial resolution and excitation laser intensity. Additionally, we noted that the fast movemnet of the clusters in the nucleus required a frame rate of ∼20 seconds to accurately capture their dynamics. These two live imaging requirements led to rapid photobleaching, within 2-3 minutes, due to the poor photostability of mVenus (***Figure 3B***).

Our two newly developed, more photostable Dorsal fusions, Dorsal-meGFP and Dorsal-mNeonGreen, enabled us to perform longer-term imaging of these Dorsal clusters. Indeed, clusters were clearly visible using both fusions (***Figure 3C,D***). Further, while the measurable fluorescence intensity was reduced to half of the starting intensity within four minutes for the Dorsal-mVenus line, the fluorescence intensity of the Dorsal-meGFP line remained above 80% of the starting intensity past six minutes (***Figure 3B***). This increased photostability enabled us to image and track Dorsal clusters for the full length of a nuclear cycle, as shown by the comparison between cluster movies of the Dorsal-mVenus and the Dorsal-meGFP fusions shown in ***Figure 3E*** and F.

With the ability to visualize Dorsal clusters in real time, we sought to uncover how they interact with target genes to regulate gene expression. To make this possible, we simultaneously imaged the Dorsal-mNeonGreen clusters and with the transcriptional activity of a *snail* reporter construct— a target of Dorsal—labeled using the MS2 system (***Bertrand et al., 1998***; ***Garcia et al., 2013***; ***Lucas et al., 2013***) over the course of an entire nuclear cycle (***Figure 4A***). Here, nascent mRNA molecules are labeled using mCherry, such that sites of nascent transcript formation are visible as fluorescent puncta.

**Figure 4.**
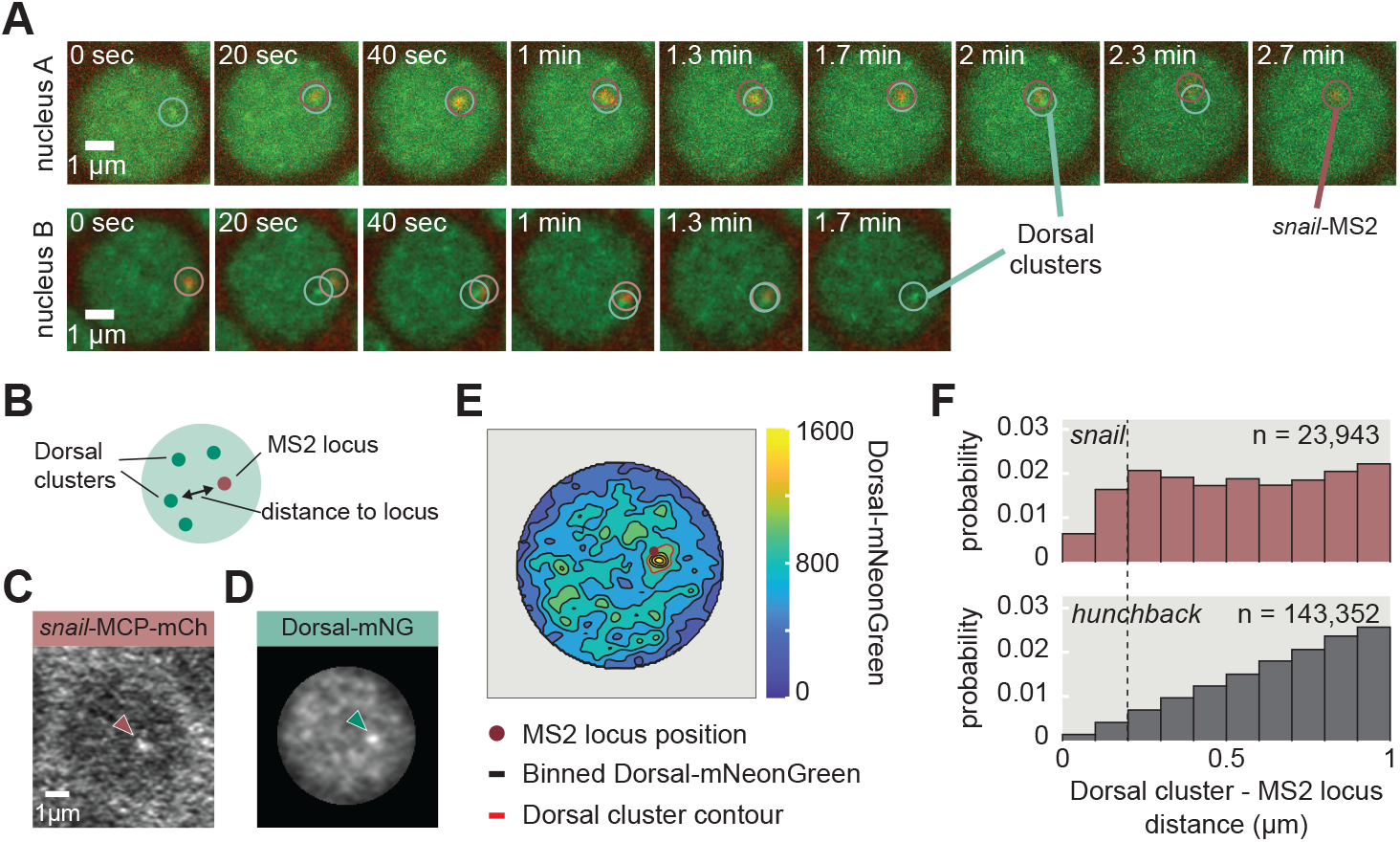
Dorsal clusters preferentially associate with transcriptional loci. **(A)** Two sample nuclei illustrating that clusters are often spatially associated with active transcriptional loci. In the top montage, nucleus A, a *snail* transcription locus turns on seemingly within a Dorsal cluster and the two remain tightly associated for approximately two minutes until the Dorsal cluster dissolves into the nuclear Dorsal background. In the bottom montage, nucleus B, a Dorsal cluster emerges from the background pool of nuclear Dorsal and moves to associate with an already-active *snail* transcription locus. The two remain associated for approximately one minute until the locus turns off, while the Dorsal cluster remains visible. **(B)** Cartoon illustrating the measurement of pairwise distances between an actively-transcribing locus labeled with MS2 and the Dorsal clusters in a nucleus. **(C-D)** Example snapshots of the same nucleus with (C) an actively-transcribing *snail* labeled by MCP-mCherry and (D) a Dorsal-mNeonGreen cluster that is nearby. **(E)** Contour level sets used to bin the Dorsal-mNeonGreen intensity are shown for the example nucleus from (D), with the second highest contour level, deemed the “Dorsal cluster” contour, outlined in red. The position of the *snail*-MS2-mCherry spot from (C) is marked by a red dot. **(F)** Distribution of pairwise distances between all Dorsal clusters in a nucleus and the *snail* (top) or and *hunchback* (bottom) locus in that same nucleus.

To investigate how the dynamics of these Dorsal clusters might dictate *snail* gene expression dynamics, we sought to characterize the positions of all clusters in a nucleus and compare them to the positions of actively transcribing *snail* loci as shown schematically in ***Figure 4B***, and as exemplified using representative images in ***Figure 4C*** and D. As the definition of a cluster is challenging due to their varying intensity and size, along with a high background intensity from nuclear Dorsal fluorescence, we adopted a segmentation approach based on contour level sets. The two highest contour heights were defined as the “cluster” levels (***Figure 4E***).

Using these contour-based cluster segmentation results, we calculated the pairwise distances between an actively transcribing, MS2-labelled locus and all the clusters detected in the same nucleus, at the same z-plane. Speficially, we quantified the distribution of pairwise cluster-locus distances for the Dorsal-target gene *snail* (***Figure 4F***). As a negative control to which to compare *snail*, we generated the same distance distribution for the non-target gene *hunchback* (***Figure 4F***).

We found that there is a population of clusters that are within 300 nm of actively transcribing *snail* loci. This population was not present near non-target *hunchback* loci. These results suggest that the population of clusters in the vicinity of the *snail* reporter is a subset of clusters that are preferentially associated with loci targeted by Dorsal. In companion papers posted alongside this work, we leverage these reagents to further explore the dynamics of these clusters and their role in regulating transcriptional activity (***Fallacaro et al., 2025***; ***Dima et al., 2025***).

### 2.5 Photoconvertible fluorescent proteins uncover Dorsal single-molecule dynamics

The recent development of lattice light sheet microscopy (***Chen et al., 2014***) has made it possible to quantify the binding dynamics of transcription factors—such as the maternally-deposited transcription factor Bicoid (***Mir et al., 2017***) and the uniformly distributed pioneer-like transcription factor Zelda (***Mir et al., 2018a***)—in living, developing fruit fly embryos. However, single molecule detection of abundant transcription factors such as Bicoid, Zelda and Dorsal on a lattice light sheet microscope requires a fluorescent protein with two key properties: high signal-to-noise ratio (SNR) and sparse labeling. For these reasons, the existing Dorsal-mVenus fusion was not a suitable choice for single-molecule measurements.

Our two new photoconvertible Dorsal fusions, Dorsal-mEos4a and Dorsal-Dendra2, solve both of these challenges, with improved SNR over Dorsal-mVenus and intrinsic photoconvertible properties, allowing for the measurement of the *in vivo* binding dynamics of individual Dorsal molecules. As a proof-of-concept, we imaged these new fusion under a custom MOSAIC microscope in the lattice light sheet imaging modality (***Chen et al., 2014***).

Excitation with a 488 nm laser enabled bulk measurements of the unconverted Dorsal-mEos4a (***Figure 5A,C***; ***Figure S5A,C***) and Dorsal-Dendra2 (***Figure S5E,G***) fusion proteins in the ventral nuclei of fruit fly embryos, where Dorsal is the most highly concentrated. Exciting the sample with the 488 nm laser also shows His2-GFP, enabling precise judgement of developmental stage. We then photoconverted a subpopulation of the Dorsal fusion proteins with a low power of a 560 nm laser and imaged the resulting photoconverted Dorsal-mEos4a (***Figure 5B***; ***Figure S5B***) and Dorsal-Dendra2 (***Figure 5F***) proteins using 560 nm excitation.

**Figure 5.**
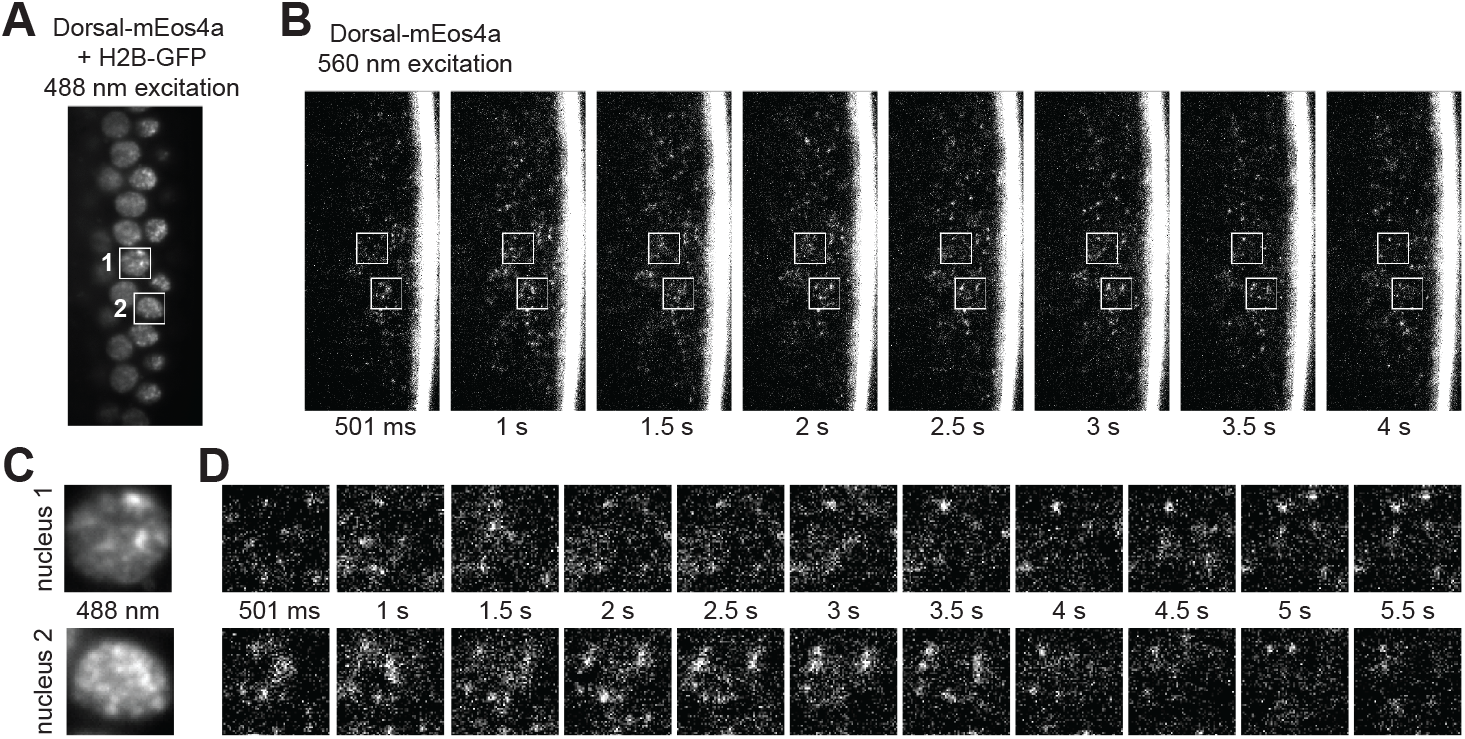
Single molecule detection of Dorsal-mEos4a fusion proteins in live embryos. **(A)** Snapshot of non-photoconverted Dorsal-mEos4a excited by a 488 nm laser line to show the location of five ventral nuclei with high nuclear Dorsal levels. Nuclei 1 and 2 are labeled with white text. **(B)** Movie stills showing a series of single-molecule detections of a photoconverted portion of Dorsal-mEos4a molecules, which were photoconverted by a 405 nm laser to be excitable by a 560 nm laser. Images were taken with a 500 ms exposure of 560 nm light approximately twice a second. **(C)** Images zooming into the nuclei marked in (A). **(D)** Image series for the two nuclei labeled in (A) single-molecule detections of photoconverted Dorsal-mEos4a.

The resulting photoconverted subpopulation of Dorsal-mEos4a was sparse enough to identify single molecules of the Dorsal fusions bound to the DNA of individual nuclei (***Figure 5D***; ***Figure S5D***). These single molecules had high signal-to-noise ratio and were photostable enough to be tracked for up to 8 s (***Figure 5D***; ***Figure S5D***). However, the photoconverted subpopulation of Dorsal-Dendra2 was initially too high, resulting in an signal-to-noise ratio that was too low for accurate single-molecule tracking (***Figure 5F***). As a result, to achieve single-molecule tracking of Dorsal-Dendra2 over the same time-scale as Dorsal-mEos4a, we first imaged under 560 nm excitation for approximately 4 min(***Figure S5F***) to bleach most of the photoconverted molecules. Only then was the unbleached, photoconverted subpopulation small enough to achieve the required signal-to-noise ratio and sparsity to track for a similar length of time (***Figure S5H***) as the Dorsal-mEos4a molecules. Thus, our new photoconvertible fluorescent Dorsal fusions are an ideal substrate to carry out single-molecule measurements of the binding dynamics of this transcription factor, as well as how these dynamics dictate output transcriptional dynamics of target genes.

To measure Dorsal binding dynamics at the single-molecule level, we performed live imaging of photoconverted Dorsal-mEos4a and His2B-mEos2.3 in Drosophila embryos using a range of exposure times (50 ms, 100 ms, and 500 ms). Due to their stable interaction with chromatin, His2B is widely seen as a benchmark for stably bound molecules (***Mir et al., 2018a***; ***Hansen et al., 2017***; ***Teves et al., 2016***). For short exposures (50 ms), traces for both stably bound and freely diffusing molecules are visible; whereas, at long exposures, the contribution of freely diffusing Dorsal molecules is blurred out, leaving only detections for molecules stably bound to DNA (***Mir et al., 2017, 2018a***). Due to their tight association with chromatin, we, therefore, expect His2B molecules to defocalize less frequently than Dorsal across all exposure times, leading to a longer average trace length for His2B. To confirm this, we constructed survival probability curves, which reflect the distribution of trajectory lengths, for Dorsal-mEos4a and His2B-mEos3.2. Indeed, for all exposure times, His2B trajectories were consistently longer than those of Dorsal (***Figure 6A***). Given that the effects of nuclear and chromatin motion, as well as photobleaching, are expected to be similar for the bound populations of both Dorsal and His2B, the longer His2B trajectories suggest two key conclusions: first, that His2B remains bound for longer durations than Dorsal, and second, that unbinding—not photobleaching—is the primary factor limiting Dorsal trajectory length. Therefore, we will utilize the 500 ms exposure data to extract the dwell time for Dorsal molecules bound to chromatin.

**Figure 6.**
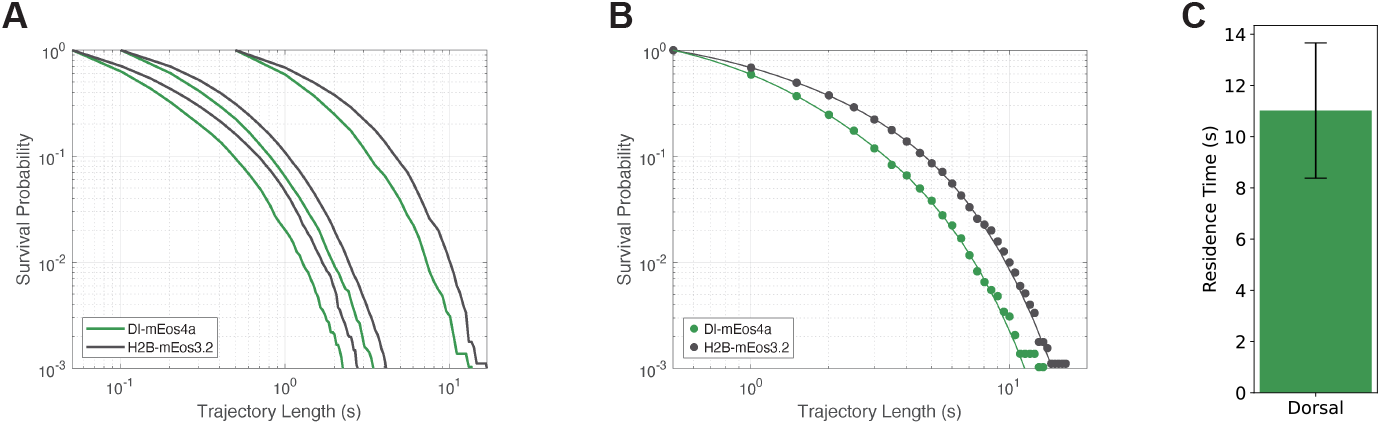
Survival Probability and Residence Time of Single Dorsal-mEos4a Proteins in Live Embryos. **(A)** Survival probability as a function of trajectory length for Dorsal (green) and Histone control (grey) at exposure times of 50 ms (left), 100 ms (center), and 500 ms (right). **(B)** Double-exponential fits of the survival probability calculated from the 500 ms exposure data. **(C)** Residence time of Dorsal-mEos4a (11.02 ± 2.64 s), estimated from the double-exponential fit.

To quantify genome average residence times, we fit the 500 ms survival probability distributions for Dorsal and His2B to a two-exponential decay model. As has been shown previously, the slow and fast time constants associated with the two exponents can be interpreted as the off-rates associated with non-specific and specific binding, respectively (***Mir et al., 2017, 2018b***; ***Hansen et al., 2017***). The resulting fit for Dorsal is then bias corrected for photobleaching and defocalization using the fit to the His2B data. This correction is based on the long-lived population of His2B associating with chromatin much longer than the dynamic range of our measurement time. The maximum trajectory lengths that we measure for His2B are thus only limited by photo-bleaching and defocalization (see Methods 4.8.3) (***Hansen et al., 2017***; ***Mir et al., 2018a***). Using this approach, we estimate an average residence time of 11 ± 2 s for Dorsal (***Figure 6C***).

These results demonstrate the utility of photoconvertible fluorescent proteins for dissecting transcription factor dynamics in vivo and are consistent with previous measurements of other transcription factors in Drosophila, such as Bicoid (***Mir et al., 2017***) and Zelda (***Mir et al., 2018a***), and in cell culture systems (***Lammers et al., 2020b***; ***Lu and Lionnet, 2021***), which have also revealed subpopulations with residence times on the order of a few seconds.

## 3 Discussion

Leveraging the advancing biochemical and optical properties of newly engineered fluorescent proteins can open new scientific avenues and help answer previously unresolved questions. However, the rate-limiting step in adopting these fluorescent proteins—particularly for essential targets like early developmental transcription factors—is achieving functional fusion with the protein of interest without disrupting its normal biological activity. In this study, we selected a panel of fluorescent proteins with useful properties for longitudinal imaging and single molecule tracking experiments, such as increased brightness, increased photostability, and the ability to photoswitch. We characterized their effects on function when fused to the maternal transcription factor Dorsal in the fruit fly embryo, and demonstrated their potential to uncover biological phenomena that were previously inaccessible.

We identified four fluorescent proteins (meGFP, mNeonGreen, mEos4a, Dendra2) that can be successfully fused to Dorsal while maintaining the ability of the Dorsal-fluorescent protein allele to function when present in two copies. However, these fusions did not fully rescue embryonic variability as homozygous alleles, let alone as heterozygotes combined with a *dorsal* mutant allele. We discovered that the DsRed marker commonly used to determine the successful CRISPR-mediated knock-in of sequences into the genome, and not the nature of the fluroescent protein or the linker between this protein and Dorsal, was the culprit behind the loss of viability of our Dorsal fusions. Indeed, both mRNA and protein levels were significantly reduced by the presence of the DsRed cassette in the 3’UTR of the *dorsal* gene. Upon removal of this cassette, both mRNA and protein levels recovered to wild-type levels and embryonic viability was restored. The mechanism by which the presence of DsRed decreases *dorsal* mRNA levels remains to be uncovered. We speculate that, due to the alteration of the 3’UTR, such reduction likely results from an alteration of the *dorsal* mRNA lifetime. Regardless, these results serve as a reminder to the fly community that, although convenient for the purposes of tracking alleles, markers such as the DsRed cassette might significantly compromise the very developmental processes we seek to characterize.

Having characterized these fusions, we demonstrated their potential for shedding light on the mechanisms of Dorsal action in two contexts. First, we showed how the increased photosability of our novel Dorsal-mNeonGreen and Dorsal-meGFP fusions made it possible to track subnuclear clusters of Dorsal protein within the nucleus. Second, we demonstrated the feasibility of using our fusions of Dorsal to photoactivatable fluorescent proteins mEos4a and Dendra2 for the characterization of Dorsal DNA binding at the single-molecule level.

For most developmental genes studied to date, the exact nature and timing of the molecular mechanisms that underlie regulation of their transcription, and the nature of the bursts by which this transcription is characterized still remain elusive (***Rodriguez and Larson, 2020***; ***Lammers et al., 2020b***; ***Leyes Porello et al., 2023***; ***Meeussen and Lenstra, 2024***). The growing body of evidence that sequence-specific and general transcription factors exist in transient clusters—also referred to as hubs, microenvironments, or condensates—of high local concentration (***Mir et al., 2017, 2018a***; ***Tsai et al., 2017***; ***Sabari et al., 2018***; ***Cho et al., 2018, 2016***; ***Zamudio et al., 2019***; ***Klosin et al., 2020***) has provided a tantalizing new way of thinking about how these molecular mechanisms could play out, as well as a potential avenue for measuring correlations with transcriptional bursting. However, so far it has been challenging to establish the functional role of these clusters in regulating transcription (***McSwiggen et al., 2019***).

Using our Dorsal fusions to photostable fluorescent proteins, we imaged its clusters in space and time and discovered that, on average, Dorsal clusters are more likely to be found in the vicinity of its target gene *snail* when compared to its non-target gene *hunchback*. In parallel collaborative work, we have used the reagents presented here to further study the interaction between Dorsal clusters and the transcriptional dynamics of Dorsal target genes (***Fallacaro et al., 2025***; ***Dima et al., 2025***). Thus, our new Dorsal-mNeonGreen and Dorsal-meGFP fusions provide an ideal tool to study the spatiotemporal underpinnings of Dorsal nuclear dynamics and the role these dynamics play in dictating transcriptional control.

Finally, our new fusions of Dorsal to photoactivatable proteins made it possible to, for the first time, detect individual molecules of this important fly transcription factor as they bind and unbind from the DNA. Our proof of concept promises to make it possible to measure the Dorsal dwell time on the DNA as it has been previously done for Bicoid and Zelda (***Mir et al., 2017, 2018a***), and to relate the localization and dynamics of this binding to the regulation of Dorsal target genes.

To sum up, our new fusions of Dorsal to novel fluorescent proteins will make it possible to reveal the mechanistic underpinnings of Dorsal and its regulation of transcription. We envision that these protein fusions will become a valuable resource for the fly community in performing the quantitative experiments necessary to reach a predictive understanding of cellular decision making in development.

## 4 Methods and Materials

### 4.1 Plasmids

To generate the Dorsal fluorescent protein knock-in alleles, we used a previously published CRISPR/Cas9 protocol (***Gratz et al., 2015***; ***Alamos et al., 2023***). The CRISPR donor plasmids were modified from a *dl-6G-mVenus-DsRed* donor plasmid (***Alamos et al., 2023***). The donor plasmid carries the following insertion sequence: a 6xGlycine (6G) linker followed by the mVenus protein coding sequence, a stop codon, a 3xP3-DsRed2-SV40polA cassette flanked by PiggyBac transposon sites, and the endogenous 3’UTR sequence. The whole insertion sequence is flanked by two ∼ 1 kb homology arms that target the endogenous Dorsal stop codon.

To generate the new fusion plasmids, the mVenus sequence was replaced with the desired fluorescent protein sequence (meGFP, mNeonGreen, mEos3.2, mEos4a, mEos4b, or Dendra2), while all other elements were left unmodified. Alternative linker plasmids were also generated for two of the fluorphore fusions, mEos3.2 and mNeonGreen. The 6G linker (GGGGGG) of the *dl-6G-mEos3.2* plasmid was replaced by three alternative linkers: 6G-10GS (GGGGGGGGGGSGSGGS), 6G-helix (GGGGGGMSKGEEL; the MSKGEEL portion is the N-terminal helix of mVenus), and LL (LongLinker, SGDSGVYKTRAQASNSAVDGTAGPGSTGSS; a gift from Michael Stadler). The 6G linker of the *dl-6G-mNeonGreen* plasmid was replaced by the LL linker. All alternative linker sequences were generated via gene synthesis by GenScript (Rijswijk, Netherlands). Cloning and sequencing to confirm the final plasmid sequence was performed by GenScript, Inc. (Rijswijk, Netherlands).

For the synthetic guide RNA (sgRNA) plasmid, we used the previously published plasmid, *pU6-DorsalgRNA1*, which expresses a synthetic guide RNA (sgRNA or gRNA) (GUUGUGAAAAAGGUAU-UACG) that targets a sequence in the C-terminus of Dorsal on Chromosome 2 (***Alamos et al., 2023***). The list of all plasmids described in the current study can be found in ***Table S2***, and full sequences for all plasmids can be accessed at https://benchling.com/garcialab/f_/THClp5A3-dorsal-fusions-manuscript/.

### 4.2 Transgenic Fly Lines

Each fluorescent protein fusion donor plasmid was co-injected with the pU6-DorsalgRNA1 plasmid into embryos expressing Cas9 under the control of *vasa* regulatory sequences in the ovary (BDSC #51324) by BestGene, Inc. (Chino Hills, CA, USA). Surviving adults were crossed to either *w1118* or *yw* stocks and their offspring were screened for DsRed fluoresence in the adult eyes. Transformants were balanced with CyO by crossing to *yw* ; *Sp/CyO* ; *+* to generate a stable lines. Several independent integrations were established as stable lines. The CRISPR insertions were confirmed by PCR using primers recognizing the left homology arm, TS-F (GAGGGCGACAAAGGCAAAGA) and Donor-R (CGCCACCACCTGTTCCTGT), and the right homology arm, Donor-F (GGGCAGCTTCACTCCTTTCT) and TS-R (TACGCCGCACTAACGAATCT). The 3xP3-DsRed2-SV40polA eye marker cassette was initially left in all lines to allow for simplified crossing schemes to other transgenic lines.

To generate the *yw; Dorsal-meGFP-DsRed, eNosx2-MCP-mCherry / CyO* ; *+* and *yw; Dorsal-meGFP-DsRed, eNosx2-MCP-mCherry / CyO* ; *+* transgenic lines were generated by recombining *yw; Dorsal-meGFP-DsRed / CyO* ; *+* or *yw; Dorsal-mNeonGreen-DsRed / CyO* ; *+* (***Table S1***) with *yw; eNosx2-MCP-mCherry @VK22 / CyO* ; *+*. Female progeny lacking the CyO balancer (i.e. curly wings) were crossed to *yw* ; *Sp/CyO* ; *+* males and the progeny of that cross were screened for individuals that had both the DsRed and white+ markers visible in the adult eye, as well as the CyO balancer (curly wings) present. Secondary, confocal microscopy screening of the embryos of the resulting stable lines was done to confirm that the lines expressed both mCherry and meGFP or mNeonGreen in the embryo.

The full list of fly lines described in this study can be found in ***Table S1***.

### 4.3 Outcrossing

*yw* ; *Dorsal-Fluorescent Protein-DsRed / CyO* ; *+* females were crossed to *yw* males. The female progeny were screened for DsRed expression and DsRed+ flies were then crossed again to *yw* males. This outcrossing was done for 6-8 generations. The final generation of female progeny was re-balanced with *CyO*, by crossing to *yw* ; *Sp/CyO* ; *+* to re-generate a stable line.

### 4.4 Embryo hatch test

To test for maternal Dorsal function in our Dorsal-FP CRISPR knock-in fusion lines, we measured embryo hatching rates for embryos laid by non-virgin females that carry two copies of the Dorsal-FP CRISPR fusion allele (i.e. homozygotes). Homozygotes were selected by screening for an absence of the balancer chromosome *CyO* and curly wings. Males were a mix of heterozygous and homozygous flies, which means that these embryo hatch tests were *not* an accurate indicator of zygotic Dorsal function, as the embryos themselves could either be heterozygous (one untagged, wildtype *dorsal* allele and one *dorsal*-FP allele) or homozygous (two *dorsal*-FP alleles). The following step-by-step protocol was used for each embryo hatch test:

1. Prepare cages with females homozygous for Dorsal-FP-DsRed and mixed homozygous and heterozygous males from the same line.
2. Transfer embryos to a juice agar plate in rows each containing 5 or 10. Record the total number of embryos mounted, aiming for at least 100 embryos per hatch test.
3. Place a small dab of yeast paste at one edge of the plate so the larvae crawl to it after they hatch.
4. Place small agar plate in a larger, covered, petri dish and leave for ∼ 36 hours at room temperature (∼ 22°*C*).
5. Count the number of empty chorions (eggshells) to determine how many embryos are “hatched”, and count the number of embryos still in their chorion to determine the number of “unhatched” embryos.
6. Confirm that the number of hatched embryos matches the difference between the number of embyros mounted and the number of unhatched embryos.

Each round of hatch tests included an agar plate of embryos from a cage of *yw* females and males and another agar plate of embryos from a cage of *yw* ; *Dorsal-6G-mVenus-DsRed ;+* females and males, both of which served as positive controls and points of comparison across different biological replicates and days.

Embryo hatch tests were performed both prior to and after outcrossing.

### 4.5 DsRed removal

DsRed marker was removed using PiggyBac transposase following an established protocol (***Nyberg and Carthew, 2022***). The fly stock having PiggyBac transposase transgene used for the crosses is w[1118]; Herm3xP3-ECFP, *α*tubuling-piggyBacK10M6 (BDSC 32070). The removal of DsRed was confirmed by checking the absence of DsRed expression in the fly eyes as well as by PCR checks using genomic DNA of each fly line as templates. To ensure the genomic edits for the FP of interest is intact, PCR checks were performed and embryos from each fly line were imaged using primers

- DsRed: ATGGCCTCCTCCGAGGACGT and CTACAGGAACAGGTGGTGGC
- mNeonGreen: GTGAGCAAGGGCGAGGAGGA and CTTGTACAGCTCGTCCATGC
- mEGFP: GTGAGCAAGGGCGAGGAGCT and CTTGTACAGCTCGTCCATGC
- Dendra2: AACACCCCGGGAATTAACCT and CCACACCTGGCTGGGCAGGG
- mEos4a: GTTAGTGCGATTAAGCCAGA and TCGTCTGGCATTGTCAGGCA

### 4.6 Quantitative real-time PCR (qPCR)

To measure the mRNA levels driven by our Dorsal fusions, embryos were collected from homozygous females of both types (with DsRed, ΔDsRed) as well as wild-type fly line *yw* as control. Grape juice plates were changed twice separated by one hour. The plates were then kept in the cages for 45 minutes before embryo collection. The embryos were washed from the plates to a mesh and dechorionated using 100% bleach for 3 minutes. The embryos were then washed with deionized water to remove residual bleach. Only the embryos younger than nc 13 were selected and washed with 500 *μ*l phosphate-buffered saline with Tween 20 in a microcentrifuge tube. RNA isolation was performed using standard TRIzol-based extraction method (Invitrogen, USA). The extracted RNA was treated with DNase I (Thermo Scientific, USA) to remove genomic DNA and purified using the Monarch RNA Cleanup Kit (New England Biolabs, USA) to remove any residual salts from the Dnase buffer. 170 ng of the collected RNA was reverse transcribed using the ProtoScript II First Strand cDNA Synthesis Kit (New England Biolabs, USA) following the standard protocol. *actin* was used as the reference gene. The primer pairs used were:

- *dl*: TGG CTT TTC GCA TCG TTT CCA G and TGT GAT GTC CAG GGT ATG ATA GCG
- *actin*: CCG TGA GAA GAT GAC CCA GAT C and TCC AGA ACG ATA CCG GTG GTA C

Genomic DNA was amplified using the same PCR primers and used as template for five 10-fold serial dilutions to generate the standard curve for the qPCR. The cDNA from fly lines with DsRed, ΔDsRed, yw were used as templates for qPCR using the *dl* and *actin* primer sets described above. All the reactions were performed in triplicates. The qPCR was performed using iTaq Universal SYBR Green Supermix (Bio-Rad, USA) in the CFX Opus 96 Real-Time PCR System (Bio-Rad, USA). The quantification cycle, *C*_*q*_ were determined using CFX Maestro Software (Bio-Rad, USA). The calibration curve was generated by plotting the average *C*_*q*_ of the triplicate vs logarithm of initial template concentration for the serial dilutions (***Figure S4***). If the initial template concentration is *N*_0_, after *C*_*q*_ cycles the concentration of the products, *N*_*x*_ will be given by

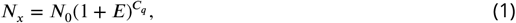

where E is the PCR efficiency. In the ideal case, *E* = 1, and the template doubles after each cycle. Taking logarithms on both sides and rearranging we get

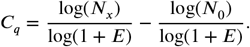

The slope of the calibration curve is therefore given by

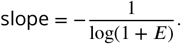

Thereby the PCR efficiency, E, of each primer pair can be determined from the slope of the calibration curve (***Bustin et al., 2009***) by calculating

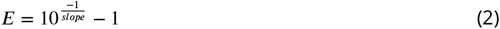

The 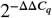 method is widely used for the calculation of the relative gene expression ratio, *R* using qPCR. Here, *C*_*q*_ represents the number of cycles required for the amplified product to reach a threshold level of *N*_*x*_ (Equation 1). In our case, this corresponds to

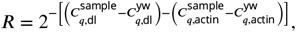

where 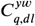 and 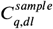 are the quantification cycle for the control line *yw* and the sample (either with DsRed or ΔDsRed) for the target gene *dl* and 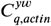 and 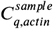 are the same for the reference gene *actin*.

However, this formula for the calculation of *R* assumes the PCR efficiency of each primer pairs to be the theoretical maximum of 1 (hence 2 corresponding to (E+1)). This can be an invalid assumption for most real-life cases. Therefore, we used a modified version that takes the experimentally determined efficiencies into account (***Pfaffl, 2001, 2004***).This leads to

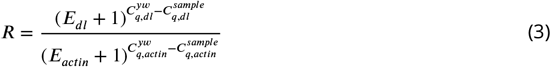

where *E*_*dl*_ and *E*_*actin*_ are the PCR efficiencies for the *dl* and *actin* primer pairs, respectively. Therefore, we calculated the relative expression ratio of *dl* based on E (equation 2) and the *C*_*q*_ deviation of the sample (either with DsRed or ΔDsRed) versus the control line *yw*, and expressed it in comparison to the reference gene *actin* (***Figure 2B***).

### 4.7 Microscopy

#### 4.7.1 Sample preparation and mounting Dorsal gradient and RICS measurements

Embryos from females having one copy (i.e. heterozygotes) of *dl-mNeonGreen-DsRed* and one copy of *His2Av-RFP* as well as embryos from females having one copy of *dl-mNeonGreen-*Δ*DsRed* and one copy of *His2Av-RFP* were collected. The *His2Av-RFP* transgene was obtained from the Bloomington Stock Center (w[*]; Pw[+mC]=His2Av-mRFP1II.2. BDSC:23651). The embryos were dechorionated in 100% bleach for 30 seconds and washed with deionized water to remove residual bleach. The embryos at nuclear cycle 10 were selected manually.

For gradient measurements, embryos were end-on mounted in 1% low melt agarose, with the anterior pole touching the coverslip. For RICS experiments, embryos were mounted on their ventral side manually.

#### 4.7.2 Cluster analysis and single-molecule experiments

Non-virgin, homozygous females and males from each Dorsal-fluorescent protein fusion line were crossed together in a cage. Virgin females from a line expressing a maternally-deposited transcription factor and the MS2 coat proteins (e.g. *yw; Dorsal-mVenus-DsRed, pNos-MCP-mCherry; Dorsal-mVenus, pNos-MCP-mCherry, His2Av-iRFP*) were crossed to males carrying an MS2 reporter reporter gene.

Cages were fed a paste of dry, activated yeast mixed with water, which was placed on a petri dish lid containing grape juice agar and changed out at least once per day. Fly cages were allowed to lay for 90 to 120 minutes prior to embryo collection. For confocal microscopy, embryos were then mounted on microscopy slides in Halocarbon 27 Oil (Sigma-Aldrich, H8773) in between a #1.5 glass coverslip and a membrane semipermeable to oxygen (Lumox film, Starstedt, Germany) as described in ***Garcia et al. (2013***) and ***Bothma et al. (2018***). For lattice lightsheet microscopy, embryos were mounted on a glue-treated 5-mm-diameter glass coverslip (Warner Instruments, no. 64-0700) and covered with phosphate-buffered saline (PBS), as described in ***Mir et al. (2017***).

#### 4.7.2 Confocal microscopy

##### Dorsal gradient and RICS experiments

Dorsal gradient measurements and RICS experiments were performed on a Zeiss LSM 900 confocal microscope. For gradient measurements, the cross-section was imaged 150 µm from the anterior pole from nuclear cycle 10 until gastrulation in 30-second time intervals with a frame time of 10 s using an LD C-Apochromat 40x/1.1 W Corr objective. Analysis of the Dorsal gradient was done following previously published protocols (***Liberman et al., 2009***; ***Reeves et al., 2012***). Briefly, nuclear segmentation was performed based on the His2Av-RFP signal. The amplitude of the gradient representing the Dorsal amount in the ventral-most nuclei and the basal level representing the non-zero amount of Dorsal in the dorsal-most nuclei were determined by Gaussian-fitting to the nuclear Dl-mNeongreen signal.

For RICS experiments, images were collected at 5x zoom (pixel size of 31.95 nm) and a frame time of 5.06 s using a C-Apochromat 40x/1.2 W autocorr objective. RICS analysis was performed following previously published protocols (***Schloop et al., 2024***; ***Al Asafen et al., 2024***; ***Dima and Reeves, 2024***).

##### General Dorsal fusion characterization

Data collection for Dorsal gradient assessment was performed on a Leica SP8 scanning confocal microscope (Leica Microsystems, Biberach, Germany). Each Dorsal-FP fusion was excited at 488 nm and its fluorescent signal detected by a Hybrid Detector (HyD) set to photon counting mode with a spectral window of 496-546 nm. Average laser powers were not quantified, but were substantially higher for mEos4a, mEos3.2, and Dendra2 as compared to meGFP and mNeonGreen to compensate for the formers’ poor quantum yield in their green states. Pinhole was set to 1.0 Airy units (AU) for an emission peak wavelength of 509 nm. All data was taken with a 63x, 1.4 NA oil objective using bidirectional scanning.

##### Cluster analysis

Data collection for the cluster analysis (***Figure 4***) results were performed on a Zeiss 980 laser scanning confocal microscope set to use the Airyscan2 (ZEISS, Jena, Germany). Dorsal-mVenus and MCP-mCherry were excited with argon ion laser lines wavelengths of 488 nm and 561 nm.

#### 4.7.3 Single-molecule tracking

Single molecule imaging was performed on a Multimodal Optical Scope with Adaptive Imaging Correction (MOSAIC) at the Advanced Bioimaging Center (ABC) at UC Berkeley. The MOSAIC was in lattice light sheet imaging mode (***Chen et al., 2014***), without adaptive optics.

An initial snapshot of non-photoconverted Dorsal-mEos4a or Dorsal-Dendra2, excited by a 488 nm laser was taken to locate the nuclei in the embryo. Single-molecules were detected by first photoconverting a fraction of the Dorsal-mEos4a or Dorsal-Dendra2 molecules with a 405 nm laser and exciting the photoconverted molecules with a 560 nm laser with a 500 ms exposure time. Light sheet excitation was conducted with a Special Optics 0.65 NA, 3.74 mm working water dipping lens. Fluorescence emissions were detected with a Zeiss 1.0 NA water-dipping objective (2.2 mm working distance) and recorded approximately twice a second for the duration of the movie with a 2x Hamamatsu Orca Flash 4.0 v3 sCMOS camera.

Additional Lattice Light Sheet Microscopy (LLSM) (***Chen et al., 2014***) experiments were performed at the Advanced Imaging Center at HHMI Janelia Research Campus. Embryos prepared as described above were placed into the LLSM bath containing room temperature PBS. The full details of the lattice light sheet microscope configuration are described previously (***Chen et al., 2014***). A custom Special Optics 0.65 NA, 3.74 mm working distance water dipping objective was used for excitation and a Nikon 1.1 NA, 2 mm working distance 25x water dipping objective (CFI Apo LWD 25XW) was used for detection. A square lattice pattern (Inner NA: 0.44; Outer NA: 0.55) was used for generating the lattice light sheet. Photoconversion was performed using a 405 nm laser line while 488 nm and 560 nm laser lines were used for imaging. Non-photoconverted Photoconverted Dl-mEos4a and His2B-mEos3.2 were imaged withe the 560 nm laser line. Emission was directed to two Hamamatsu Orca Flash 4.0 sCMOS cameras (Dichroic: Semrock FF560-FDi01-25×36; Camera 1 – Semrock BLP01-532R-25, Semrock NF03-488E-25, Semrock NF03-561E-25; Camera 2 – Semrock BLP01-488r-25, Semrock FF01-520/35-25). The net system magnification is 63x for a pixel size of 104 nm. For all experiments, a field of view of 608 × 256 pixels was used.

Custom scripts for the LLSM were developed for the photoconversion and single molecule imaging experiments. For both the Dorsal-mEos4a experiment and the H2B-mEos3.2 control experiment, an initial image of the histone channel was acquired (488 nm excitation, 100 ms exposure time, 23 *μ*W; note that all powers were measured entering the back focal plane of the excitation objective). For the Dorsal-mEos4a experiments, a 405 nm light sheet was scanned 10 *μ*m in 100 nm steps (101 images total) around the image plane (100 ms exposure time, 12.7 *μ*W). Due to the higher labeling density of the H2B-mEos3.2, no initial photoconversion was necessary but instead photobleaching was required to obtain an appropriate number of localizations per frame. To do so, a 560 nm light sheet was scanned 5 *μ*m in 50 nm steps (101 images total) around the image plane (100 ms exposure time, 748 *μ*W). This was repeated continuously for 30 iterations. All subsequent steps were identical for both the Dorsal-mEos4a experiments and H2B-mEos3.2 control experiments. A single plane was imaged using excitation via a 560 nm light sheet. Separate experiments were performed with 3 different exposure times (50 ms, 100 ms, and 500 ms); a total of 2000, 1000, and 300 images were collected with excitation powers of 690 *μ*W, 455 *μ*W, and 153 *μ*W, respectively. Due to the readout speed of the camera, the actual frame rate for each experiment was 19.51 Hz, 9.88 Hz, and 2.00 Hz, respectively. After the single-plane, single-molecule imaging was completed, a final image of the histone channel was collected (100 ms exposure time, 113 *μ*W) to account for any nuclear movement and assess changes in the stage of development. No deskewing was necessary for the LLSM experiments as a single plane was imaged rather than a 3D stack.

### 4.8 Image Analysis

Image processing and extraction of MS2 movies was performed in MATLAB using the custom pipeline described in ***Garcia et al. (2013***) and ***Lammers et al. (2020a***), which can be found in the public mRNADynamics Github repository. Transcription spots and nuclei were segmented with the aid of the Trainable Weka Segmentation plugin for FIJI (***Witten et al., 2011***; ***Arganda-Carreras et al., 2017***).

#### 4.8.1 Bleaching quantification

Both the Dorsal-meGFP and Dorsal-mVenus curves in ***Figure 3B*** were quantified by generating summed z-projections at each time point and calculating the fluorescence per pixel (summed fluorescence intensity in the 2D z-projection divided by the total number of pixels) for each time point. All time points were then normalized to the first time point by dividing by the total fluorescence intensity per pixel at t=0.

#### 4.8.2 Single molecule detection and tracking

We used the quot single-molecule detection package to identify and track photoconverted Dorsal-mEos4a and His2B-mEos2.3 molecules in live Drosophila embryos. Initial spot detection was performed using a log-likelihood ratio (LLR) test, which compares the likelihood of a Gaussian spot centered in a local window to that of a flat background with Gaussian-distributed noise. Detection parameters were empirically optimized for our imaging conditions: Gaussian kernel amplitude, window size, and detection threshold. For each detected spot, subpixel localization was performed using a two-step procedure. First, a radial symmetry method provided an initial estimate of the center. This was followed by maximum likelihood estimation of a symmetric, 2D integrated Gaussian point spread function (PSF) using a Levenberg-Marquardt optimization routine.

Molecule trajectories were reconstructed using a conservative tracking algorithm. Localizations were assigned to existing trajectories only if the correspondence was unambiguous—specifically, if exactly one trajectory and one localization fell within the predefined search radius. Ambiguous assignments were disallowed to avoid incorrect linking: unmatched trajectories were terminated, and unmatched localizations were treated as the start of new trajectories. Tracking parameters were adjusted for each dataset to optimize trajectory lengths while minimizing erroneous links.

#### 4.8.3 Calculation of residence times from long exposure single-molecule trajectories

Imaging with sufficiently long exposure times effectively blurs out fast-moving molecules into the background while molecules stably bound to chromatin for a significant duration of the exposure time are captured as diffraction-limited spots (***Mir et al., 2018a***; ***Hansen et al., 2017***; ***Teves et al., 2016***). Therefore, the trajectories from the 500 ms datasets are used to infer the genome average long-lived (specific) binding times.

To infer the residence time, the length of trajectories in time is used to calculate a survival probability (SP) curve (1 - cumulative distribution function of trajectory lengths). Since the SP curve contains contributions from non-specific interactions, slowly moving molecules, and localization errors a double-exponential function of the form *SP* (*t*) = *F* exp(−*k*_*ns*_*t*) + (1 − *F*) exp(−*k*_*s*_*t*) is fit to the SP curve, where *k*_*ns*_ is the off-rate for the short-lived (non-specific) interactions and *k*_*s*_ corresponds to the off-rate of long lived (specific) interactions (***Mir et al., 2018a, 2017***; ***Hansen et al., 2017***; ***Teves et al., 2016***) (6). For fitting purposes, probabilities below 10^−3^ are not used to avoid fitting the data poor tails of the distribution.

Next, since the inferred ks as described above is biased by photobleaching, and nuclear and chromatin movement, bias correction is performed using the His2B data as *k*_*s*,true_ = *k*_*s*_ −*k*_bias_, where kbias is the slower rate from the double-exponent fit to the His2B SP curve as described previously (Teves et al., 2016; Hansen et al., 2017; Chen et al., 2014a; Chen et al., 2014b). This correction is based on the assumption that photobleaching, unbinding, and loss of trajectories from motion are all independent Poisson processes. The genome wide specific residence time is then calculated as 1/ks,true. The effectiveness of this bias correction is checked by calculating the residence time from both the 100 ms and 500 ms frame rate data and observing convergence to within 1 s (Figure 4—figure supplement 2C).

## 5 Supplementary Material

Supplementary Material, including figures and tables, can be found at the end of this document.

## 6 Acknowledgments

We would like to acknowledge the contribution of Simon Alamos and Emma Luu for help with reagent preparation and fly husbandry. We also thank Thomas Graham, Mustafa Mir, Xavier Darzacq, Mike Eisen, Jenna Haines, Mike Stadler and Jacques Bothma for helpful discussions. We thank Holly Aaron for her microscopy training, advice, and support for these experiments using the Zeiss LSM 980 with Airyscan2 confocal at the CRL Molecular Imaging Center at Berkeley, RRID:SCR_017852, supported by NIH S10OD025063. We would also like to thank Gokul Upadhyayula and Gaoxiang Liu from the the Advanced Bioimaging Center (ABC) at UC Berkeley. The ABC is supported by the Chan Zuckerberg Initiative, HHMI, and the Philomathia Foundation. Further,we thank Chad Hob-son, Teng-Leong Chew, Jesse Aaron and Rachel Lee from the Advanced Imaging Center at Janelia Research Campus. The Advanced Imaging Center is supported by the Howard Hughes Medical Institute and by the Gordon and Betty Moore Foundation. MAT was supported by the National Science Foundations Graduate Research Fellowship Program (NSF GRFP). HGG was supported by NIH R01 Awards R01GM139913 and R01GM152815, by the Koret-UC Berkeley-Tel Aviv University Initiative in Computational Biology and Bioinformatics, by a Winkler Scholar Faculty Award, and by the Chan Zuckerberg Initiative Grant CZIF2024-010479. H.G.G. is also a Chan Zuckerberg Biohub–San Francisco Investigator.

## 7 Author contributions

Conceptualization: MAT, HGG

Methodology: MAT, BM, NG, NL, HGG, SD, GR

Resources: MAT, GM, HGG, SD, GR

Investigation: MAT, BM, NG, NL, HGG, SD, GR

Visualization: MAT, NG, HGG, SD

Funding acquisition: HGG, GR

Project administration: HGG

Supervision: HGG, GR

Writing – original draft: MAT

Writing – review & editing: MAT, HGG, BM, NG, SD, GR

## 9 Data and materials availability

All materials are available upon request. All data in the main text or supplementary materials are available upon request.

## Supplementary Information

### S1 Eliminating off-target CRISPR/Cas9 mutations does not improve viability

We sought to remove any off-target mutations at other genomic loci by crossing the four new Dorsal fusion lines with *yw* ; *+* ; *+* flies for 6–8 generations, in a process called out-crossing or chromosome cleaning (*Methods* ***Section 4.3***). We then conducted an embryo hatch test using homozygous females from these out-crossed lines, comparing the percentage of embryos laid after 36 hours to the original CRISPR alleles. As shown in ***Figure S1***, only one of the *dl-6G-meGFP-DsRed* lines saw an increase in their hatch rates after outcrossing, but this improvement was only modest (from 5% to 8%) and did not improve embryo viability enough to enable maintenance of a stable population. These results suggested that off-target CRISPR mutations were not a significant cause of lowered viability in our new Dorsal fusion lines.

### S2 Modifying the linker sequence does not improve viability

We also attempted to determine if the linker sequence was impacting the viability of our new fusion lines. When designing protein fusions, the length, flexibility, and composition of the linker placed between the protein of interest and fluorescent protein can all be critical to maintaining endogenous activity and function (***Chen et al., 2013***). We replaced the 6G linker (used by both ***Reeves et al. (2012***) and ***Alamos et al. (2023***)) in the *dl-6G-mNeonGreen-DsRed* plasmid with another linker, LL (literally, LongLinker; SGDSGVYKTRAQASNSAVDGTAGPGSTGSS, a kind gift of Michael Stadler). However, only 1% of embryos from females homozygous for the modified linker *dl-LL-mNeonGreen-DsRed* allele hatched, and only after outcrossing (***Figure S1***).

Additionally, in an attempt to generate a mEos3.2 fusion allele that produces any viable progeny, we replaced the 6G linker in the *dl-6G-mEos3.2-DsRed* plasmid with three other linkers: 6G-10GS (GGGGGGGGGGSGSGGS), LL, and 6G-helix (GGGGGGMSKGEEL, where MSKGEEL is the N-terminal helix of mVenus). No viable progeny were produced by females homozygous for any of these three alternative linker mEos3.2 alleles. Thus, the choice of linker did not have a stronger effect than the choice of fluoroscent protein.

### S3 Quantification of nuclear Dorsal-mNeonGreen absolute concentration

We measured the absolute concentration of nuclear Dorsal-mNeonGreen fusion proteins in the ventral nuclei of *DsRed* and Δ*DsRed* embryos using Raster Image Correlation Spectroscopy (RICS), which can quantify the concentration of fluorescent protein molecules in live cells (***Digman et al., 2005a***,b; ***Digman and Gratton, 2009***; ***Schloop et al., 2024***; ***Brown et al., 2008***; ***Al Asafen et al., 2024***) (***Section 4***). The nuclei on the ventral side of the embryos were imaged from nuclear cycle 10 until gastrulation.

RICS analysis showed that the nuclear Dorsal-mNeonGreen concentration in the *DsRed* embryos was lower than the concentration in the Δ*DsRed* across all nuclear cycles (***Figure S2A***). At the concentration maxima in nuclear cycles 13 and 14, Dorsal-mNeonGreen concentration in *DsRed* embryos was only 36% and 47% of the concentrations measured in the nuclei of Δ*DsRed* embryos, respectively (***Figure S2B***). The protein level is not reduced exactly to half as reported by our qPCR measurements (***Figure 2A***). This discrepancy might result from errors in the estimation of concentration in RICS, due to variation in the mRNA degradation rate in the two lines, or due to differences in translation efficiency of the *dorsal* mRNA.

**Figure S1.**
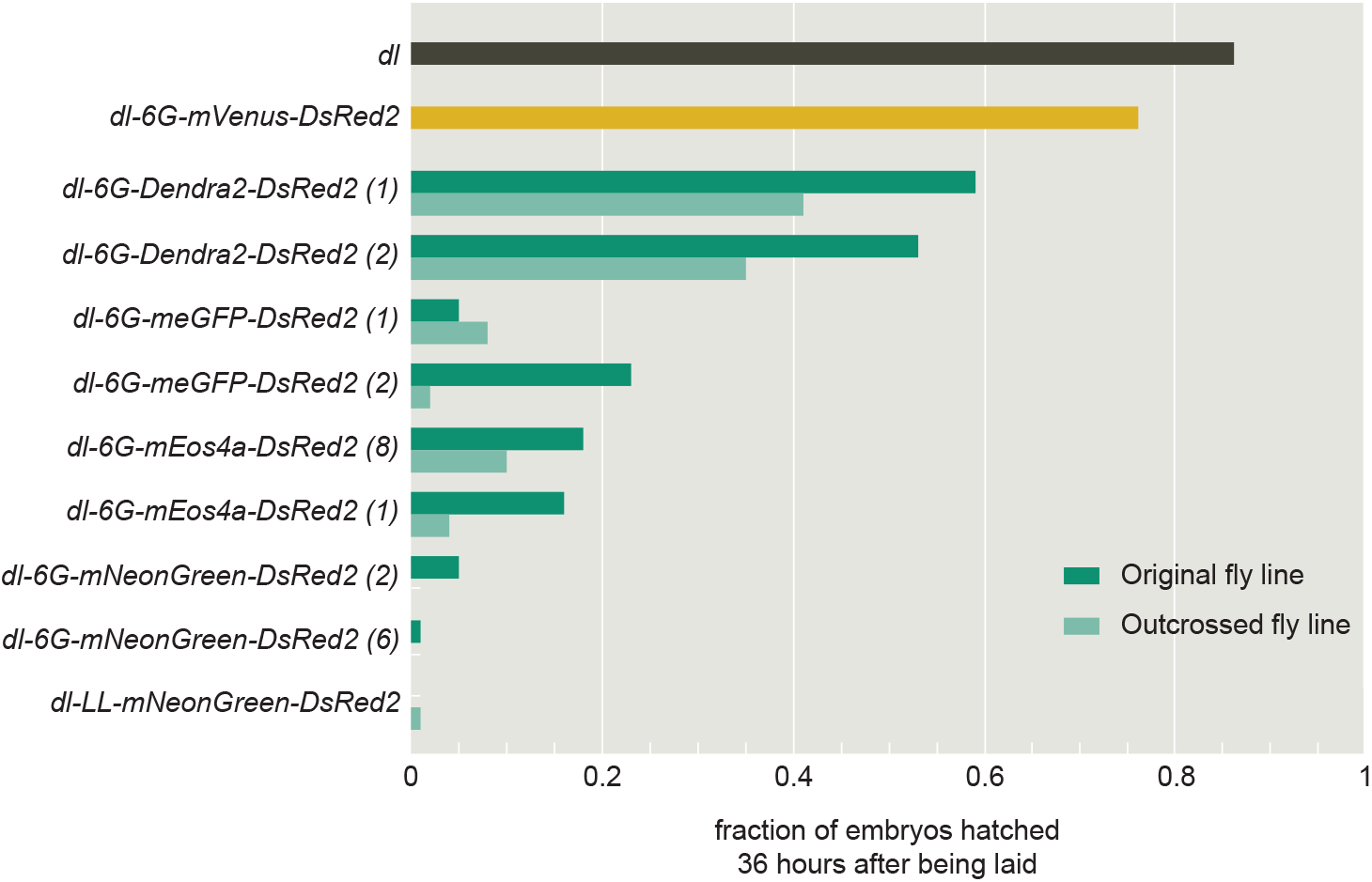
Embryo hatch test results before and after outcrossing. Hatch rates for embryos laid by females homozygous for the Dorsal fusion alleles, quantified as fraction of embryos that had hatched 36 hours after being laid. For the *dl-6G-Dendra2-DsRed2, dl-6G-meGFP-DsRed2, dl-6G-mEos4a-DsRed2*, and *dl-6G-mNeonGreen-DsRed2* alleles, homozygous females from original (top, dark bar in each pair) and outcrossed (bottom, light bar in each pair) versions of each fly line are compared. Embryo hatch rates for an untagged *dl* allele (from *yw* flies) served as a control.

**Figure S2.**
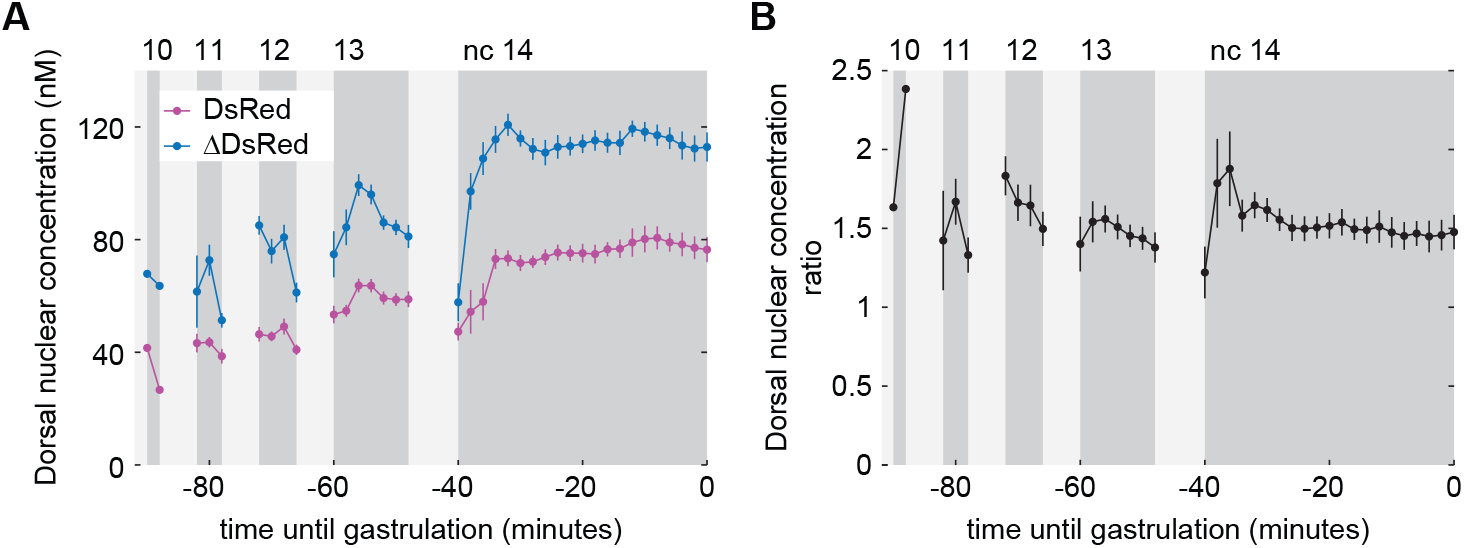
Quantification of nuclear Dorsal-mNeonGreen absolute concentration. **(A)** Absolute Dorsal-mNeonGreen protein concentration, as quantified by RICS, in the ventral nuclei of DsRed (pink) and ΔDsRed (blue) embryos during nuclear cycles 10 to 14. Error bars are the standard error of the mean. **(B)** Ratio of absolute Dorsal-mNeonGreen protein concentration in ΔDsRed to DsRed embryos in ventral nuclei. DsRed: *dl-6G-mNeonGreen-DsRed* allele; ΔDsRed: *dl-6G-mNeonGreen-*Δ*DsRed* allele.

**Figure S3.**
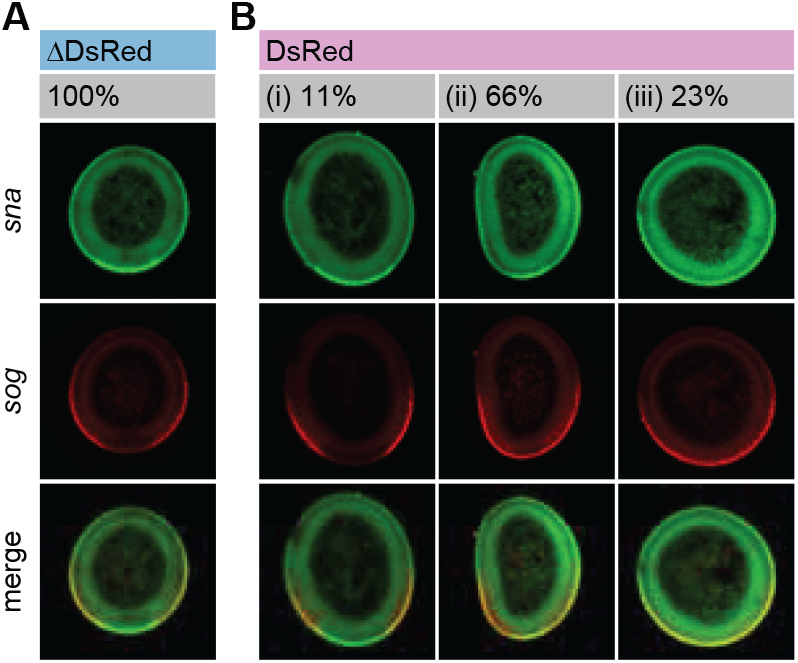
Gene expression of Dorsal-activated genes in DsRed and ΔDsRed embryos. **(A)** Gene expression of two Dorsal-activated genes, *snail* (*sna*; green) and *short gastrulation* (*sog*; red), in the cross-section of a representative *dl-6G-mNeonGreen-*Δ*DsRed* embryo, as measured by fluorescence *in situ* hybridized (FISH). 24 embryos were imaged in total. *sna* is a Type I Dorsal-regulated gene, which is activated only by the highest levels of Dorsal on the ventral side of the embryo. *short gastrulation* (*sog*) is a Type III Dorsal-regulated gene, which is activated by even the lower levels of Dorsal present on the lateral sides of the embryo (***Reeves and Stathopoulos, 2009***) and repressed by high levels of Snail protein on the ventral side of the embryo. These expression patterns are similar to those seen in wild-type embryos (***Leptin, 1991***; ***Francois et al., 1994***; ***Srinivasan et al., 2002***). **(B)** Gene expression of the two Dorsal-activated genes, *sna* (green) and *sog* (red), in the cross-section of *dl-6G-mNeonGreen-DsRed* embryos, as measured by fluorescence *in situ* hybridized (FISH). 44 embryos were imaged in total. Representative images are shown for each of the three distinct types of gene expression patterns observed: (i) 11% (5/44) of embryos exhibit normal expression of both *sna* and *sog*; (ii) 66% (29/44) of embryos exhibit a narrow *sna* pattern and, in turn, an extended *sog* pattern; and (iii) 23% (10/44) of embryos exhibit an absent *sna* pattern and, in turn, an extended *sog* pattern.

**Figure S4.**
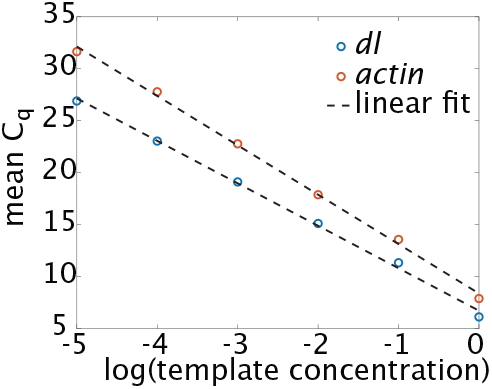
Calibration curve for the determination of PCR efficiencies of the primer pairs. Blue circles represent the average of triplicate experiments for *dl* and orange circles the same for *actin*. The slope of the linear fit is used for the calculation of PCR efficiencies of the corresponding primer pairs.

**Figure S5.**
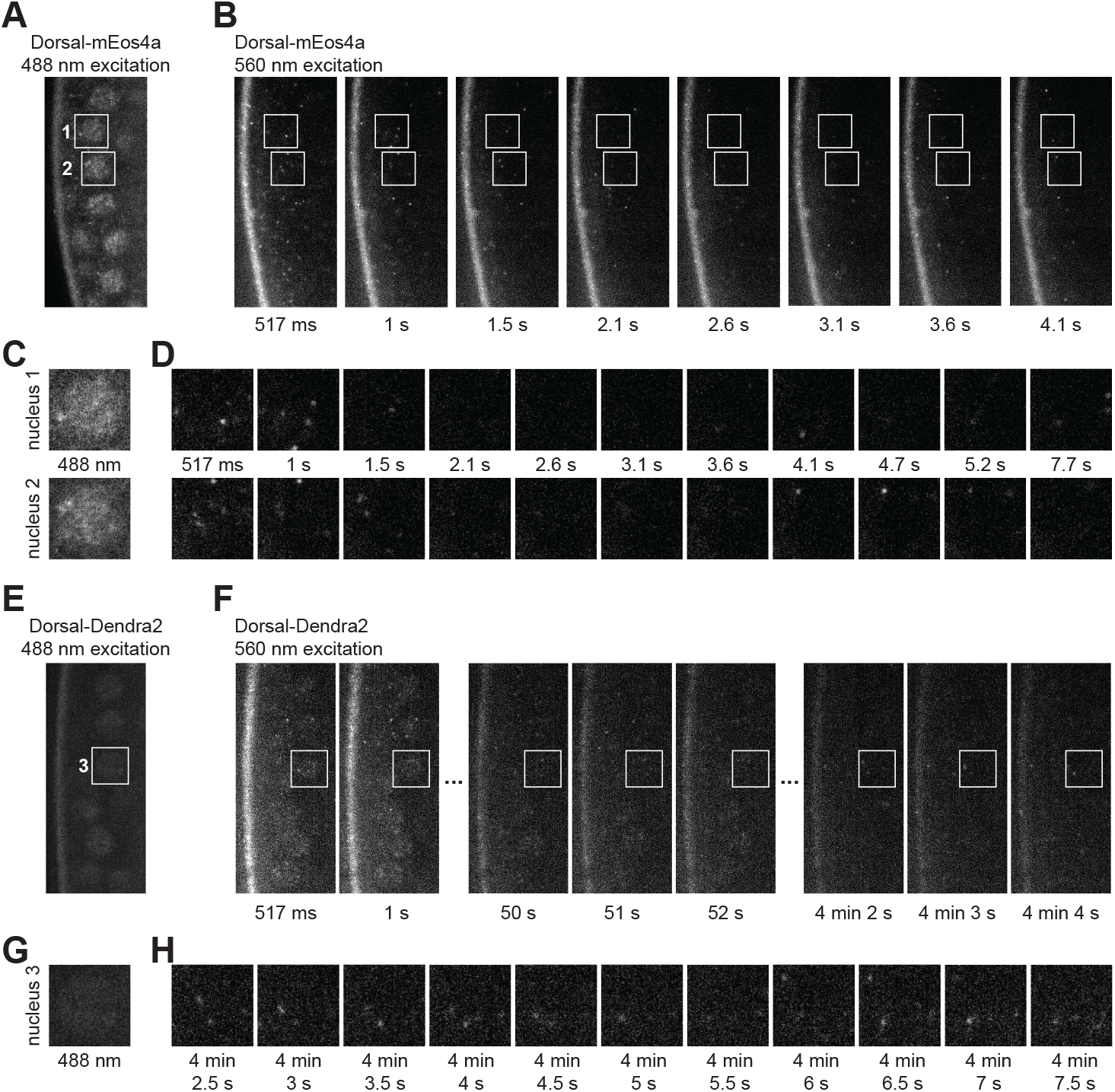
Single molecule detection of Dorsal-mEos4a and Dorsal-Dendra2 fusion proteins in live embryos. Single-molecule detection in an embryo expressing **(A-D)** Dorsal-mEos4a and **(E-H)** Dorsal-Dendra2. **(A)** Snapshot of non-photoconverted Dorsal-mEos4a excited by a 488 nm laser line to show the location of five ventral nuclei with high nuclear Dorsal levels. Nuclei 1 and 2 are labeled with white text. **(B)** Movie stills showing a series of single-molecule detections of a photoconverted portion of Dorsal-mEos4a molecules, which were photoconverted by a 405 nm laser to be excitable by a 560 nm laser. Images were taken with a 500 ms exposure of 560 nm light approximately twice a second. **(C-D)** Image series for the two nuclei labeled in (A), nucleus 1 (top) and nucleus 2 (bottom), showing **(D)** single-molecule detections of photoconverted Dorsal-mEos4a within the boundaries of each nucleus. **(E-H)** Single-molecule detection in an embryo expressing Dorsal-Dendra2. **(E)** Snapshot of non-photoconverted Dorsal-Dendra2 excited by a 488 nm laser line to show the location of five ventral nuclei with high nuclear Dorsal levels. Nucleus 3 is labeled with white text. **(F)** Movie stills showing a series of single-molecule detections of a photoconverted portion of Dorsal-Dendra2 molecules, which were photoconverted by a 405 nm laser to be excitable by a 560 nm laser. Images were taken with a 500 ms exposure of 560 nm light approximately twice a second. **(G-H)** Image series for the nucleus 3, labeled in (E), showing **(H)** single-molecule detections of photoconverted Dorsal-Dendra2 specifically within the boundaries of the nucleus.

**Table S1.** List of fly lines used in the current study.

| Transgenic Fly Lines |  |
| --- | --- |
| Genotype | Source |
| *w[1118]; +; PBacy[+mDint2] GFP[E.3xP3]=vas-Cas9VK00027* | BDSC #51324 |
| *w[1118]* | BestGene, Inc. |
| *yw* | BestGene, Inc. |
| *yw; Sp/CyO; +* | lab stock |
| *yw / w; Dorsal-6G-mVenus-DsRed; +* | Alamos et al. (2023) |
| *yw / w; Dorsal-6G-meGFP-DsRed / CyO; +* | current study |
| *yw / w; Dorsal-6G-mNeonGreen-DsRed / CyO; +* | current study |
| *yw / w; Dorsal-6G-Dendra2-DsRed / CyO; +* | current study |
| *yw / w; Dorsal-6G-mEos4a-DsRed / CyO; +* | current study |
| *yw / w; Dorsal-6G-mEos4b-DsRed / CyO; +* | current study |
| *yw / w; Dorsal-6G-mEos3.2-DsRed / CyO; +* | current study |
| *yw / w; Dorsal-10GS-mEos3.2-DsRed / CyO; +* | current study |
| *yw / w; Dorsal-6G-helix-mEos3.2-DsRed / CyO; +* | current study |
| *yw / w; Dorsal-LongLinker-mEos3.2-DsRed / CyO; +* | current study |
| *yw / w; Dorsal-LongLinker-mNeonGreen-DsRed / CyO; +* | current study |
| *yw; Dorsal-mVenus-DsRed, pNos-MCP-mCherry; Dorsal-mVenus, pNos-MCP-mCherry, His2Av-iRFP* | Alamos et al. (2023) |
| *yw; eNosx2-MCP-mCherry / CyO; +* | current study |
| *yw; Dorsal-meGFP-DsRed, eNosx2-MCP-mCherry / CyO; +* | current study |
| *yw; Dorsal-mNeonGreen-DsRed, eNosx2-MCP-mCherry / CyO; +* | current study |
| *w; snaBAC-MS2; +* | Bothma et al. (2015) |
| *yw; hbP2P-MS2; +* | Garcia et al. (2013) |

**Table S2.** List of plasmids described in this study. Full sequences for all plasmids introduced in the current study can be accessed through a Benchling repository at https://benchling.com/garcialab/f_/THClp5A3-dorsal-fusions-manuscript/

| Plasmids |  |
| --- | --- |
| Name | Source |
| pU6-DorsalgRNA1 | *Alamos et al. (2023)* |
| *dl-6G-mVenus-DsRed* | *Alamos et al. (2023)* |
| *dl-6G-meGFP-DsRed* | current study |
| *dl-6G-mNeonGreen-DsRed* | current study |
| *dl-6G-Dendra2-DsRed* | current study |
| *dl-6G-mEos4a-DsRed* | current study |
| *dl-6G-mEos4b-DsRed* | current study |
| *dl-6G-mEos3.2-DsRed* | current study |
| *dl-10GS-mEos3.2-DsRed* | current study |
| *dl-6G-helix-mEos3.2-DsRed* | current study |
| *dl-LongLinker-mEos3.2-DsRed* | current study |
| *dl-LongLinker-mNeonGreen-DsRed* | current study |
| *snailBAC/MS2-yellow (snaBAC)* | *Bothma et al. (2015)* |
| *pIB-hbP2P-24xMS2v5-lacZ-tub3'UTR* | *Garcia et al. (2013)* |

## References

Al Asafen, H., Clark, N., Goyal, E., Jacobsen, T., Siddika Dima, S., Chen, H.-Y., Sozzani, R., and Reeves, G. (2024). Dorsal/NF-B exhibits a dorsal-to-ventral mobility gradient in the Drosophila embryo.

Alamos, S., Reimer, A., Westrum, C., Turner, M. A., Talledo, P., Zhao, J., Luu, E., and Garcia, H. G. (2023). Minimal synthetic enhancers reveal control of the probability of transcriptional engagement and its timing by a morphogen gradient. Cell Systems, 14(3).

Andreassi, C. and Riccio, A. (2009). To localize or not to localize: mRNA fate is in 3UTR ends.

Arganda-Carreras, I., Kaynig, V., Rueden, C., Eliceiri, K. W., Schindelin, J., Cardona, A., and Sebastian Seung, H. (2017). Trainable Weka Segmentation: a machine learning tool for microscopy pixel classification. Bioinformatics, 33(15):2424–2426.

Bertrand, E., Chartrand, P., Schaefer, M., Shenoy, S. M., Singer, R. H., and Long, R. M. (1998). Localization of ASH1 mRNA particles in living yeast. Mol Cell, 2(4):437–45.

Boka, A. P., Mukherjee, A., and Mir, M. (2021). Single-molecule tracking technologies for quantifying the dynamics of gene regulation in cells, tissue and embryos. Development, 148(18).

Bothma, J. P., Garcia, H. G., Ng, S., Perry, M. W., Gregor, T., and Levine, M. (2015). Enhancer additivity and non-additivity are determined by enhancer strength in the Drosophila embryo. eLife, 4.

Bothma, J. P., Norstad, M. R., Alamos, S., and Garcia, H. G. (2018). LlamaTags: A Versatile Tool to Image Transcription Factor Dynamics in Live Embryos. Cell.

Brown, C., Dalal, R., Hebert, B., Digman, M., Horwitz, A., and Gratton, E. (2008). Raster image correlation spectroscopy (RICS) for measuring fast protein dynamics and concentrations with a commercial laser scanning confocal microscope. J Microsc, 229:78–91.

Bustin, S., Benes, V., Garson, J., Hellemans, J., Huggett, J., Kubista, M., Mueller, R., Nolan, T., Pfaffl, M., and Shipley, G. (2009). The MIQE guidelines: Minimum information for publication of quantitative real-time PCR experiments. Clin Chem, 55:611–622.

Buxbaum, A., Haimovich, G., and Singer, R. (2015). In the right place at the right time: Visualizing and understanding mRNA localization.

Chen, B.-C., Legant, W. R., Wang, K., Shao, L., Milkie, D. E., Davidson, M. W., Janetopoulos, C., Wu, X. S., Hammer, J. A., Liu, Z., English, B. P., Mimori-Kiyosue, Y., Romero, D. P., Ritter, A. T., Lippincott-Schwartz, J., Fritz-Laylin, L., Mullins, R. D., Mitchell, D. M., Bembenek, J. N., Reymann, A.-C., Bohme, R., Grill, S. W., Wang, J. T., Seydoux, G., Tulu, U. S., Kiehart, D. P., and Betzig, E. (2014). Lattice light-sheet microscopy: Imaging molecules to embryos at high spatiotemporal resolution. Science, 346(6208):1257998–1257998.

Chen, S. X., Osipovich, A. B., Ustione, A., Potter, L. A., Hipkens, S., Gangula, R., Yuan, W., Piston, D. W., and Magnuson, M. A. (2011). Quantification of factors influencing fluorescent protein expression using rmce to generate an allelic series in the rosa26 locus in mice. Dis Model Mech, 4(4):537–47.

Chen, X., Zaro, J. L., and Shen, W.-C. (2013). Fusion protein linkers: Property, design and functionality. Advanced Drug Delivery Reviews, 65(10):1357–1369.

Cho, W.-K., Jayanth, N., English, B. P., Inoue, T., Andrews, J. O., Conway, W., Grimm, J. B., Spille, J.-H., Lavis, L. D., Lionnet, T., and Cisse, I. I. (2016). RNA Polymerase II cluster dynamics predict mRNA output in living cells. eLife, 5.

Cho, W.-K., Spille, J.-H., Hecht, M., Lee, C., Li, C., Grube, V., and Cisse, I. I. (2018). Mediator and RNA polymerase II clusters associate in transcription-dependent condensates. Science, page eaar4199.

Cormack, B. P., Valdivia, R. H., and Falkow, S. (1996). FACS-optimized mutants of the green fluorescent protein (GFP). Gene, 173(1):33–38.

Cranfill, P. J., Sell, B. R., Baird, M. A., Allen, J. R., Lavagnino, Z., de Gruiter, H. M., Kremers, G. J., Davidson, M. W., Ustione, A., and Piston, D. W. (2016). Quantitative assessment of fluorescent proteins. Nat Methods, 13(7):557– 62.

Digman, M., Brown, C., Sengupta, P., Wiseman, P., Horwitz, A., and Gratton, E. (2005a). Measuring fast dynamics in solutions and cells with a laser scanning microscope. Biophys J, 89:1317–1327.

Digman, M. and Gratton, E. (2009). Analysis of diffusion and binding in cells using the RICS approach. Microsc Res Tech, 72:323–332.

Digman, M., Sengupta, P., Wiseman, P., Brown, C., Horwitz, A., and Gratton, E. (2005b). Fluctuation correlation spectroscopy with a laser-scanning microscope: Exploiting the hidden time structure. Biophys J, 88:33– 36.

Dima, S. and Reeves, G. (2024). Bulk-level maps of pioneer factor binding dynamics during Drosophila maternal-to-zygotic transition.

Dima, S. S., Turner, M. A., Garcia, H. G., and Reeves, G. T. (2025). Global maps of transcription factor properties reveal threshold-based formation of DNA-bound and mobile clusters. bioRxiv, page 2025.04.24.650477.

Donovan, B. T., Huynh, A., Ball, D. A., Patel, H. P., Poirier, M. G., Larson, D. R., Ferguson, M. L., and Lenstra, T. L. (2019). Live-cell imaging reveals the interplay between transcription factors, nucleosomes, and bursting. The EMBO Journal, 38(12).

Fallacaro, S., Mukherjee, A., Turner, M. A., Garcia, H. G., and Mir, M. (2025). Transcription factor hubs exhibit gene-specific properties that tune expression. bioRxiv, page 2025.04.07.647578.

Francois, V., Solloway, M., O’Neill, J. W., Emery, J., and Bier, E. (1994). Dorsal-ventral patterning of the Drosophila embryo depends on a putative negative growth factor encoded by the short gastrulation gene. Genes & Development, 8(21):2602–2616.

Garcia, H. G., Tikhonov, M., Lin, A., and Gregor, T. (2013). Quantitative imaging of transcription in living Drosophila embryos links polymerase activity to patterning. Curr Biol, 23(21):2140–5.

Gilmore, T. D. (2006). Introduction to NF-B: players, pathways, perspectives. Oncogene, 25(51):6680–6684.

Gratz, S. J., Rubinstein, C. D., Harrison, M. M., Wildonger, J., and O’Connor-Giles, K. M. (2015). CRISPR-Cas9 Genome Editing in Drosophila. Current Protocols in Molecular Biology, 111(1).

Gurskaya, N. G., Verkhusha, V. V., Shcheglov, A. S., Staroverov, D. B., Chepurnykh, T. V., Fradkov, A. F., Lukyanov, S., and Lukyanov, K. A. (2006). Engineering of a monomeric green-to-red photoactivatable fluorescent protein induced by blue light. Nature Biotechnology, 24(4):461–465.

Hansen, A. S., Pustova, I., Cattoglio, C., Tjian, R., and Darzacq, X. (2017). Ctcf and cohesin regulate chromatin loop stability with distinct dynamics. Elife, 6.

Hong, J.-W., Hendrix, D. A., Papatsenko, D., and Levine, M. S. (2008). How the Dorsal gradient works: Insights from postgenome technologies. Proceedings of the National Academy of Sciences, 105(51):20072–20076.

Kawasaki, K. and Fukaya, T. (2023). Functional coordination between transcription factor clustering and gene activity. Mol Cell, 83(10):1605–1622 e9.

Klosin, A., Oltsch, F., Harmon, T., Honigmann, A., Jülicher, F., Hyman, A. A., and Zechner, C. (2020). Phase separation provides a mechanism to reduce noise in cells. Science, 367(6476):464–468.

Kopek, B. G., Paez-Segala, M. G., Shtengel, G., Sochacki, K. A., Sun, M. G., Wang, Y., Xu, C. S., van Engelenburg, S. B., Taraska, J. W., Looger, L. L., and Hess, H. F. (2017). Diverse protocols for correlative super-resolution fluorescence imaging and electron microscopy of chemically fixed samples. Nature Protocols, 12(5):916–946.

Kuersten, S. and Goodwin, E. (2003). The power of the 3 UTR: Translational control and development.

Lammers, N. C., Galstyan, V., Reimer, A., Medin, S. A., Wiggins, C. H., and Garcia, H. G. (2020a). Multimodal transcriptional control of pattern formation in embryonic development. Proc Natl Acad Sci U S A, 117(2):836– 847.

Lammers, N. C., Kim, Y. J., Zhao, J., and Garcia, H. G. (2020b). A matter of time: Using dynamics and theory to uncover mechanisms of transcriptional bursting. Curr Opin Cell Biol, 67:147–157.

Leptin, M. (1991). twist and snail as positive and negative regulators during Drosophila mesoderm development. Genes & Development, 5(9):1568–1576.

Leyes Porello, E. A., Trudeau, R. T., and Lim, B. (2023). Transcriptional bursting: stochasticity in deterministic development. Development, 150(12).

Liberman, L. M., Reeves, G. T., and Stathopoulos, A. (2009). Quantitative imaging of the Dorsal nuclear gradient reveals limitations to threshold-dependent patterning in Drosophila. Proceedings of the National Academy of Sciences, 106(52):22317–22322.

Lu, F. and Lionnet, T. (2021). Transcription factor dynamics. Cold Spring Harb Perspect Biol, 13(11).

Lucas, T., Ferraro, T., Roelens, B., De Las Heras Chanes, J., Walczak, A. M., Coppey, M., and Dostatni, N. (2013). Live imaging of Bicoid-dependent transcription in Drosophila embryos. Curr Biol, 23(21):2135–9.

Mayr, C. (2019). What are 3 utrs doing? Cold Spring Harb Perspect Biol 11.

McKinney, S. A., Murphy, C. S., Hazelwood, K. L., Davidson, M. W., and Looger, L. L. (2009). A bright and photo-stable photoconvertible fluorescent protein. Nature Methods, 6(2):131–133.

McSwiggen, D. T., Mir, M., Darzacq, X., and Tjian, R. (2019). Evaluating phase separation in live cells: diagnosis, caveats, and functional consequences. Genes & Development, page genesdev;gad.331520.119v1.

Meeussen, J. V. W. and Lenstra, T. L. (2024). Time will tell: comparing timescales to gain insight into transcriptional bursting. Trends Genet, 40(2):160–174.

Mir, M., Reimer, A., Haines, J. E., Li, X.-Y., Stadler, M., Garcia, H., Eisen, M. B., and Darzacq, X. (2017). Dense Bicoid hubs accentuate binding along the morphogen gradient. Genes & Development, (31):1784–1794.

Mir, M., Stadler, M. R., Ortiz, S. A., Hannon, C. E., Harrison, M. M., Darzacq, X., and Eisen, M. B. (2018a). Dynamic multifactor hubs interact transiently with sites of active transcription in Drosophila embryos. eLife, page 27.

Mir, M., Stadler, M. R., Ortiz, S. A., Hannon, C. E., Harrison, M. M., Darzacq, X., and Eisen, M. B. (2018b). Dynamic multifactor hubs interact transiently with sites of active transcription in drosophila embryos. Elife, 7:e40497.

Nyberg, K. and Carthew, R. (2022). CRISPR-/Cas9-Mediated Precise and Efficient Genome Editing in Drosophila. In Methods in Molecular Biology, pages 135–156. Humana Press Inc.

Paez-Segala, M. G., Sun, M. G., Shtengel, G., Viswanathan, S., Baird, M. A., Macklin, J. J., Patel, R., Allen, J. R., Howe, E. S., Piszczek, G., Hess, H. F., Davidson, M. W., Wang, Y., and Looger, L. L. (2015). Fixation-resistant photoactivatable fluorescent proteins for CLEM. Nature Methods, 12(3):215–218.

Pfaffl, M. (2001). A new mathematical model for relative quantification in real-time RT–PCR. Nucleic Acids Res, 29:45– 45.

Pfaffl, M. (2004). Quantification strategies in real-time PCR. AZ of quantitative PCR, 1:89–113.

Pimmett, V. L., McGehee, J., Trullo, A., Douaihy, M., Radulescu, O., Stathopoulos, A., and Lagha, M. (2024). Opto-genetic manipulation of nuclear Dorsal reveals temporal requirements and consequences for transcription. bioRxiv: The Preprint Server for Biology, page 2024.11.28.623729.

Reeves, G., Trisnadi, N., Truong, T., Nahmad, M., Katz, S., and Stathopoulos, A. (2012). Dorsal-Ventral Gene Expression in the Drosophila Embryo Reflects the Dynamics and Precision of the Dorsal Nuclear Gradient. Developmental Cell, 22(3):544–557.

Reeves, G. T. and Stathopoulos, A. (2009). Graded Dorsal and Differential Gene Regulation in the Drosophila Embryo. Cold Spring Harbor Perspectives in Biology, 1(4):a000836.–a000836. tex.ids: Reeves2009a.

Rodriguez, E. A., Campbell, R. E., Lin, J. Y., Lin, M. Z., Miyawaki, A., Palmer, A. E., Shu, X., Zhang, J., and Tsien, R. Y. (2017). The Growing and Glowing Toolbox of Fluorescent and Photoactive Proteins. Trends in Biochemical Sciences, 42(2):111–129.

Rodriguez, J. and Larson, D. R. (2020). Transcription in living cells: Molecular mechanisms of bursting. Annu Rev Biochem, 89:189–212.

Sabari, B. R., Dall’Agnese, A., Boija, A., Klein, I. A., Coffey, E. L., Shrinivas, K., Abraham, B. J., Hannett, N. M., Zamudio, A. V., Manteiga, J. C., Li, C. H., Guo, Y. E., Day, D. S., Schuijers, J., Vasile, E., Malik, S., Hnisz, D., Lee, T. I., Cisse, I. I., Roeder, R. G., Sharp, P. A., Chakraborty, A. K., and Young, R. A. (2018). Coactivator condensation at super-enhancers links phase separation and gene control. Science, 361(6400):eaar3958.

Schloop, A., Hiremath, S., Shaikh, R., Williams, C., and Reeves, G. (2024). Spatiotemporal dynamics of NF-B/Dorsal inhibitor IB/Cactus in Drosophila blastoderm embryos.

Singh, A. P., Wu, P., Ryabichko, S., Raimundo, J., Swan, M., Wieschaus, E., Gregor, T., and Toettcher, J. E. (2022). Optogenetic control of the Bicoid morphogen reveals fast and slow modes of gap gene regulation. Cell Rep, 38(12):110543.

Srinivasan, S., Rashka, K. E., and Bier, E. (2002). Creation of a Sog Morphogen Gradient in the Drosophila Embryo. Developmental Cell, 2(1):91–101.

Szostak, E. and Gebauer, F. (2013). Translational control by 3’-UTR-binding proteins. Brief Funct Genomics, 12:58–65.

Teves, S. S., An, L., Hansen, A. S., Xie, L., Darzacq, X., and Tijan, R. (2016). A dynamic mode of mitotic bookmarking by transcription factors. Elife, (5):e22280.

Tsai, A., Muthusamy, A. K., Alves, M. R., Lavis, L. D., Singer, R. H., Stern, D. L., and Crocker, J. (2017). Nuclear microenvironments modulate transcription from low-affinity enhancers. eLife, 6.

Wagh, K., Stavreva, D. A., Upadhyaya, A., and Hager, G. L. (2023). Transcription factor dynamics: One molecule at a time. Annu Rev Cell Dev Biol, 39:277–305.

Wei, M. T., Chang, Y. C., Shimobayashi, S. F., Shin, Y., Strom, A. R., and Brangwynne, C. P. (2020). Nucleated transcriptional condensates amplify gene expression. Nat Cell Biol, 22(10):1187–1196.

Witten, I. H., Frank, E., and Hall, M. A. (2011). Data Mining: Practical Machine Learning Tools and Techniques. Elsevier.

Yamada, S., Whitney, P. H., Huang, S. K., Eck, E. C., Garcia, H. G., and Rushlow, C. A. (2019). The Drosophila Pioneer Factor Zelda Modulates the Nuclear Microenvironment of a Dorsal Target Enhancer to Potentiate Transcriptional Output. Curr Biol, 29(8):1387–1393 e5.

Zacharias, D. A., Violin, J. D., Newton, A. C., and Tsien, R. Y. (2002). Partitioning of Lipid-Modified Monomeric GFPs into Membrane Microdomains of Live Cells. Science, 296(5569):913–916.

Zamudio, A. V., Dall’Agnese, A., Henninger, J. E., Manteiga, J. C., Afeyan, L. K., Hannett, N. M., Coffey, E. L., Li, C. H., Oksuz, O., Sabari, B. R., Boija, A., Klein, I. A., Hawken, S. W., Spille, J.-H., Decker, T.-M., Cisse, I. I., Abraham, B. J., Lee, T. I., Taatjes, D. J., Schuijers, J., and Young, R. A. (2019). Mediator Condensates Localize Signaling Factors to Key Cell Identity Genes. Molecular Cell, 76(5):753–766.e6.

Zhang, M., Chang, H., Zhang, Y., Yu, J., Wu, L., Ji, W., Chen, J., Liu, B., Lu, J., Liu, Y., Zhang, J., Xu, P., and Xu, T. (2012). Rational design of true monomeric and bright photoactivatable fluorescent proteins. Nature Methods, 9(7):727–729.

